# Acquired temozolomide resistance instructs patterns of glioblastoma behavior in gelatin hydrogels

**DOI:** 10.1101/2023.11.14.567115

**Authors:** Victoria Kriuchkovskaia, Ela K. Eames, Rebecca B. Riggins, Brendan A.C. Harley

## Abstract

Acquired drug resistance in glioblastoma (GBM) presents a major clinical challenge and is a key factor contributing to abysmal prognosis, with less than 15 months median overall survival. Aggressive chemotherapy with the frontline therapeutic, temozolomide (TMZ), ultimately fails to kill residual highly invasive tumor cells after surgical resection and radiotherapy. Here, we report a three-dimensional (3D) engineered model of acquired TMZ resistance using two isogenically-matched sets of GBM cell lines encapsulated in gelatin methacrylol hydrogels. We benchmark response of TMZ-resistant vs. TMZ-sensitive GBM cell lines within the gelatin-based extracellular matrix platform and further validate drug response at physiologically relevant TMZ concentrations. We show changes in drug sensitivity, cell invasion, and matrix-remodeling cytokine production as the result of acquired TMZ resistance. This platform lays the foundation for future investigations targeting key elements of the GBM tumor microenvironment to combat GBM’s devastating impact by advancing our understanding of GBM progression and treatment response to guide the development of novel treatment strategies.

**Teaser:** A hydrogel model to investigate the impact of acquired drug resistance on functional response in glioblastoma.

## Introduction

Glioblastoma (GBM) is the most common and lethal form of primary brain cancer (1). Despite an aggressive treatment strategy, including surgical resection followed by radiation and chemotherapy, GBM has one of the highest mortality rates among all human cancers with a median survival of only 12–15 months post diagnosis (1, 2). Temozolomide (TMZ; Temodar®) remains the standard-of-care therapeutic against GBM since its FDA approval for newly diagnosed glioblastomas in 2005 (3, 4). However, this GBM treatment strategy has not led to a profound advancement in care, with TMZ providing only minimal benefit to patients (5). As a prodrug, TMZ undergoes intracellular hydrolysis into a highly potent alkylating metabolite, 3-methyl-(triazen-1-yl)imidazole-4-carboximide (MTIC). MTIC methylates purine bases of DNA (O6-guanine; N7-guanine and N3-adenine), eventually leading to apoptosis (6). However, the primary cytotoxic lesion, O6-methylguanine, can be directly repaired by a suicide DNA repair enzyme, methylguanine methyltransferase (MGMT) in GBM tumors expressing MGMT (MGMT+ tumors, over 60% of all glioblastomas) (7). Unfortunately, even patients with GBM tumors not expressing MGMT (MGMT-tumors) can suffer from other mechanisms of resistance or ultimately acquire resistance to TMZ during standard-of-care treatment (8).

The unique and complex tumor microenvironment (TME) is believed to play a key role in GBM progression (9). While it rarely metastasizes, GBM invades diffusely throughout the brain (10). The composition and cell-mediated remodeling of the TME extracellular matrix (ECM) can affect patterns of diffuse invasion and drug response (11, 12). For instance, the perivascular niche (PVN) is believed to influence patterns of GBM invasion and TMZ resistance (13). Despite *in vivo* animal models being the gold standard of preclinical drug studies, systematic examination of the influence of the matrix environment on functional changes in GBM tumor behavior is largely intractable *in vivo* (14–16). As a result, there is a significant opportunity to develop *in vitro* models to evaluate GBM drug response that can consider the role of the matrix environment and multicellular interactions on cell response. Such tools would be a valuable complement to the cell monolayer two-dimensional (2D) cultures, which offer simple and reproducible systems to evaluate drug response, but which do not provide an avenue to examine the role of the matrix environment on GBM progression. Three-dimensional (3D) cultures have recently become attractive models to recapitulate key elements of tumor pathology *in vitro* (17–22). Our lab has recently described gelatin-based hydrogel platforms to investigate pathophysiological processes of GBM progression. Notably, we showed that engineered hydrogels that contain gradients of biophysical and metabolic (e.g., brain-mimetic hyaluronic acid, hypoxia) properties induce functional shifts in GBM cell motility and therapeutic response (11, 23–25). More recently, we established an engineered model of the PVN to show that the perivascular environment promotes GBM invasion and resistance to TMZ (13, 26, 27). Separately, Tiek et al. established a robust 2D model of acquired TMZ resistance using a unique set of isogenically-matched, TMZ-sensitive vs. TMZ-resistant GBM cell lines (8MGBA vs. 8MGBA-TMZres; 42MGBA vs. 42MBGA-TMZres). Continual exposure of the parental lines, 8MGBA (derived from a frontal lobe tumor resected from a 54-year-old female) and 42MGBA (derived from a temporal lobe tumor resected from a 63-year-old male), to TMZ established TMZ resistant (TMZres) cell lines that showed increased MGMT expression and no longer underwent TMZ-induced apoptosis (28).

In this study, we combine these technologies to benchmark patterns of invasion and functional response of the isogenically-matched TMZ-sensitive vs. TMZ-resistant GBM cell lines (8MGBA vs. 8MGBA-TMZres; 42MGBA vs. 42MBGA-TMZres) in 3D gelatin hydrogels. We assess the response of the GBM cells with acquired TMZ resistance (vs. isogenically-matched TMZ-sensitive, WT cells) to physiologically relevant concentrations of TMZ in an engineered ECM gelatin platform. We report changes in drug response, cell motility (invasion), and cytokine production in response to acquired TMZ resistance. This work defines a landscape for future projects to systematically and accessibly assess the impact of key GBM-specific tumor microenvironment components on drug response and GBM progression

## Results

### Acquired Temozolomide Resistance Can Be Robustly Defined in Gelatin Hydrogels

First, we determined drug response metrics of the temozolomide (TMZ) sensitive (WT) vs. resistant (TMZres) glioblastoma (GBM) cell lines within engineered extracellular matrix (ECM) hydrogels (Fig. 1, Table 1). After encapsulating GBM cells into methacrylamide-functionalized gelatin (GelMA) hydrogels and treating cell-laden hydrogels with TMZ, we measured the effect of single-dose TMZ treatments on cell viability with alamarBlue assay™. Next, we determined growth rate (GR) inhibition metrics induced by TMZ. We observed that both TMZ resistant cell lines (8MGBA-TMZres and 42MGBA-TMZres) displayed robust drug resistance patterns when cultured within GelMA hydrogels (Fig. 1B-C). The TMZ resistant cell lines (8MGBA-TMZres and 42MGBA-TMZres) displayed significant increases in GR50 values versus WT (TMZ-sensitive) control cell lines (8MGBA-WT and 42MGBA-WT) for up to 7 days after treatment. At day 7 post treatment, the change in GR50 value was >250 µM for the 8MGBA pair (WT: 27.52 µM to TMZres: 279.43 µM; Table 1 and Fig. 1B) and >1000 µM for the 42MGBA pair (WT: 2.95 µM to TMZres: 1224.71 µM; Table 1 and Fig. 1C). Similar trends in TMZ response of the 42MGBA pair were seen at earlier (Day 3, 5) timepoints (Table 1, Fig. S1B, Fig. S2B). At day 3, 8MGBA-TMZres responded more favorably to TMZ compared to its parental cell line, 8MGBA (Fig. S1A); however, at day 5, 8MGBA-TMZres started showing robust resistance to TMZ (Fig S2A). Overall, the 42MGBA-TMZres cells displayed the greatest degree of resistance, with virtually unaffected growth rates unless treated with supraphysiological (>300 µM) TMZ concentrations (Fig. 1C).

**Fig. 1.**
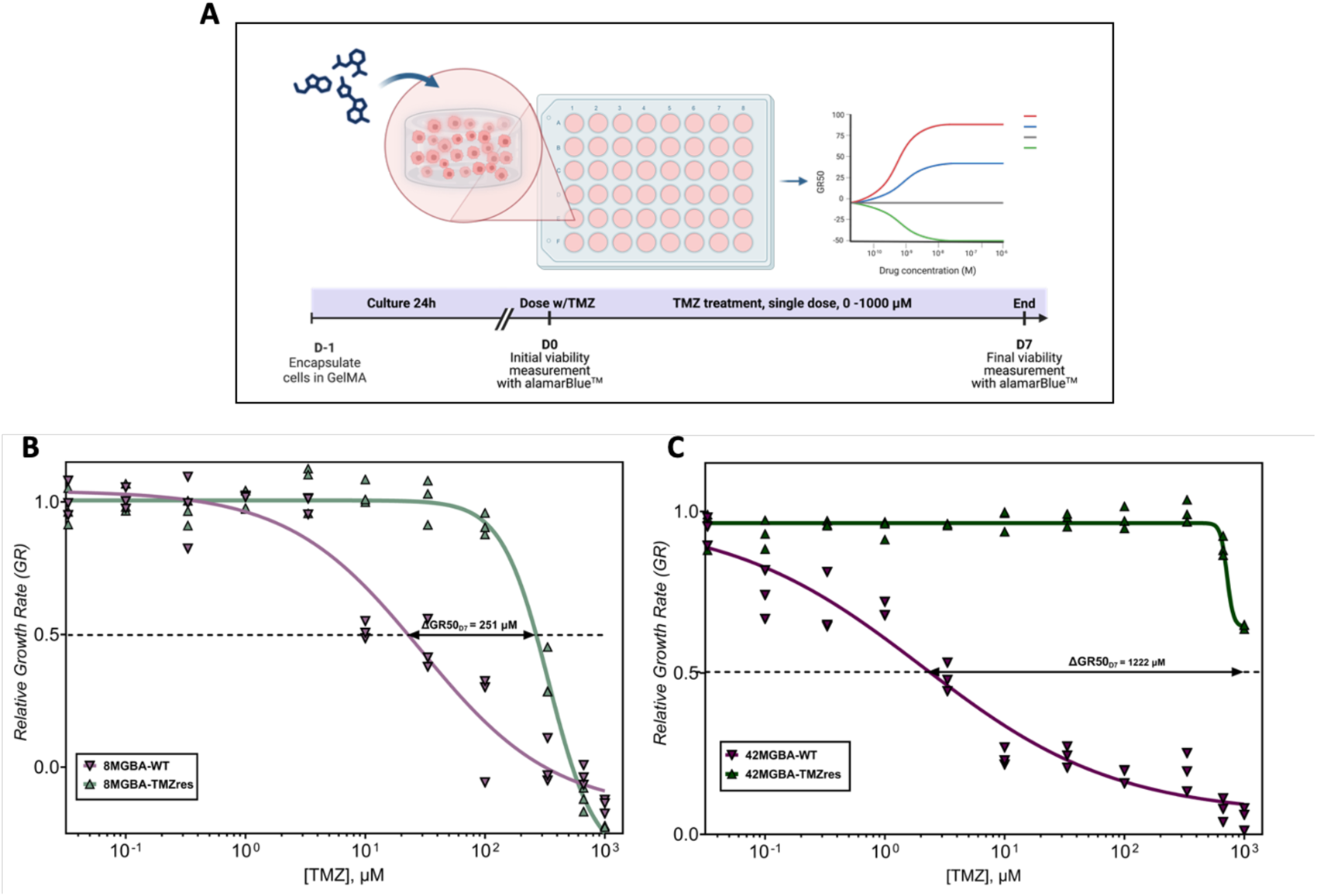
Dose-response curves of TMZ-sensitive vs. TMZ-resistant glioblastoma cell lines cultured in GelMA hydrogels. **(A)** Experimental setup to determine drug response metrics in temozolomide (TMZ) resistant vs. sensitive cell lines. First, cells were encapsulated in GelMA hydrogels. After 24 hours of culture in GelMA hydrogels, initial metabolic activity of the encapsulated cells was measured with the alamarBlue™ reagent, and then hydrogels were immediately treated with TMZ (D0). TMZ was applied to hydrogels as a single dose (without replenishing the drug) in serial dilutions of 0 – 1000 µM concentrations, with 0 µM being DMSO vehicle control. TMZ-treated hydrogels were further cultured for 7 days. Cell viability measurements were taken 3, 5 and 7 days after TMZ treatment. **(B)** Dose response, growth-rate inhibition curves of the 8MGBA vs. 8MGBA-TMZres cell lines 7 days after TMZ treatment. **(C)** Dose response, growth-rate inhibition curves of the 42MGBA vs. 42MGBA-TMZres cell lines 7 days after TMZ treatment. Each individual data point along with fitted GR curves are shown. N = 3 hydrogels per condition.

**Table 1.** Dose-response metrics of TMZ-sensitive vs. TMZ-resistant glioblastoma cell lines cultured in GelMA hydrogels. Dose response metrics, 50% inhibition of growth rate (GR50) and half maximal effective concertation (GEC50), for each TMZ-sensitive/resistant glioblastoma cell line 3, 5, and 7 days post TMZ treatment.

| *Cell Line* | *Day 3* |  | *Day 5* |  | *Day 7* |  |
| --- | --- | --- | --- | --- | --- | --- |
| | GR50, μM | GEC50, μM | GR50, μM | GEC50, μM | GR50, μM | GEC50, μM |
| 8MGBA-WT | 150.50 | 599.52 | 68.23 | 300.76 | 27.52 | 41.57 |
| 8MGBA-TMZ<sup>res</sup> | 76.08 | 248.38 | 242.88 | 316.94 | 279.43 | 383.83 |
| 42MGBA-WT | 5.84 | 393.69 | 1.29 | 1.89 | 2.95 | 2.27 |
| 42MGBA-TMZ<sup>res</sup> | 799.67 | 91930.14 | 921.41 | 676.77 | 1224.71 | 1052.17 |

### Temozolomide-Resistant GBM Cells Display Potent Resistance across a Broad Range of Physiological TMZ Doses

We subsequently compared the ability to resolve cytotoxic effects of repeated, low-concentration (metronomic) temozolomide doses on the wild type vs. TMZ-resistant 8MGBA and 42MGBA cell lines (Fig. 2A), quantifying the relative metabolic activity (fold change over DMSO vehicle control) for the 8MGBA vs. 8MGBA-TMZres cell lines (Fig. 2B-C) and the 42MGBA vs. 42MGBA-TMZres cell lines (Fig. 2D-E). Broadly, these data suggested that while WT GBM cells are responsive to single and repeated TMZ doses in gelatin hydrogels, TMZ-resistant lines show robust resistance to both single and metronomic doses (Fig. 2). Further, in some cases, repeated low-dose TMZ exposure promotes GBM cell viability.

**Fig. 2.**
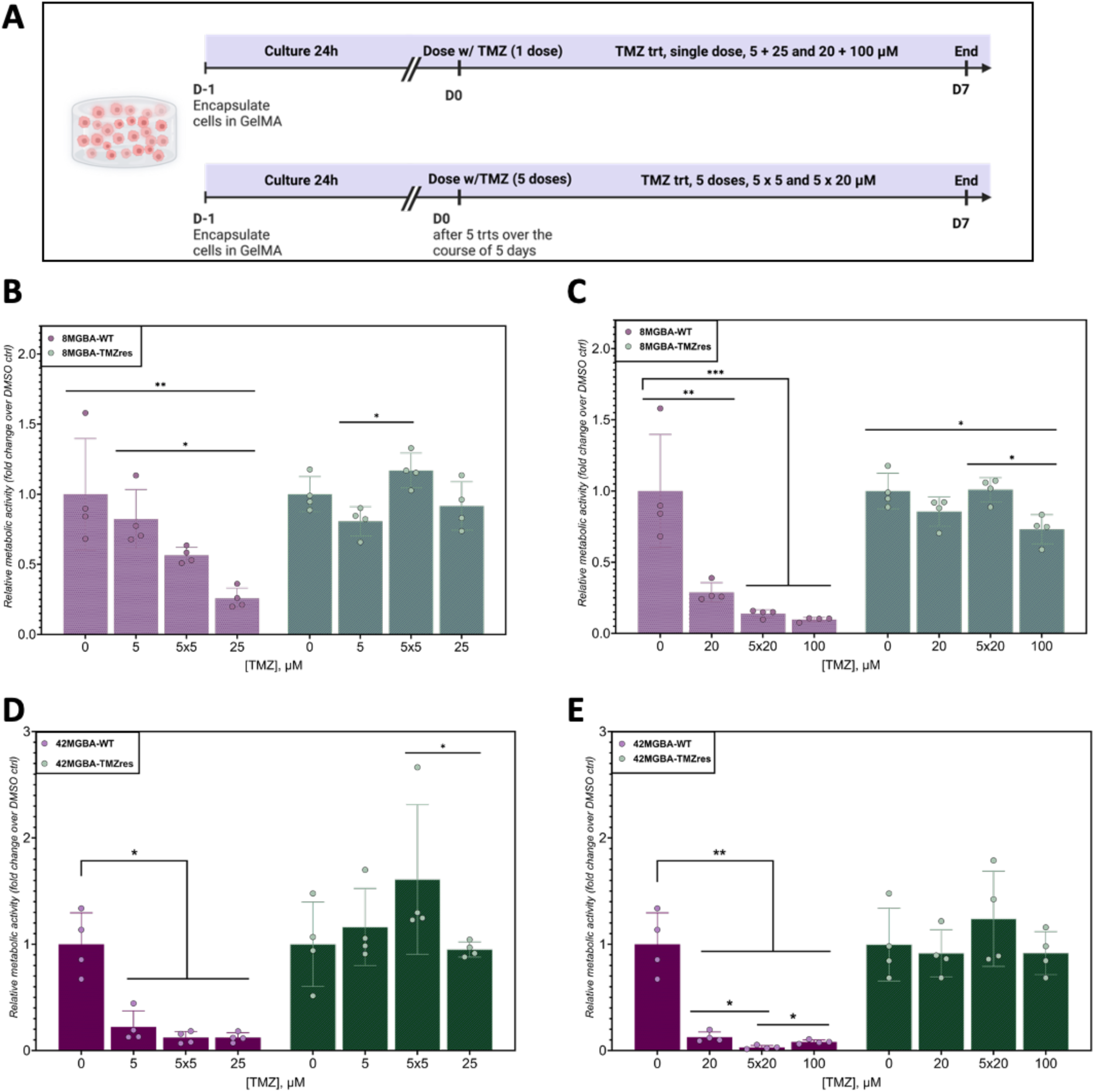
Validating drug response of TMZ-sensitive/resistant glioblastoma model at physiologically relevant drug concentrations. **(A)** Experimental setup to determine drug response to low single vs. repeated TMZ doses. First, cells were encapsulated in GelMA hydrogels. After 24 hours of culture, cell-laden GelMA hydrogels were treated with the following sets of doses: (1) a single dose of 5 µM, or 5 repeated doses of 5 µM, or a cumulative single dose of 25 µM, and similarly, (2) a single dose of 20 µM, or 5 repeated doses of 20 µM, or a cumulative single dose of 100 µM. The repeated doses were administered in daily intervals (24 hours between each dose). Cell viability was measured at days 0, 3, 5, and 7 after either the single TMZ treatment for single dose exposures or after the final TMZ treatment for repeated dose exposure. Day 0 was set to be as the day of the final TMZ treatment for each experiment. Data was normalized to day 0 measurements prior to the analysis. **(B)** Relative metabolic activity of 8MGBA-WT and 8MGBA-TMZres cells treated with a single dose of 5 µM, 5 repeated doses of 5 µM, or a cumulative single dose of 25 µM. **(C)** Relative metabolic activity of 8MGBA-WTand 8MGBA-TMZres cells treated with a single dose of 20 µM, 5 repeated doses of 20 µM, or a cumulative single dose of 100 µM. **(D)** Relative metabolic activity of 42MGBA-WT and 42MGBA-TMZres cells treated with a single dose of 5 µM, 5 repeated doses of 5 µM, or a cumulative single dose of 25 µM. **(E)** Relative metabolic activity of 42MGBA-WT and 42MGBA-TMZres cells treated with a single dose of 20 µM, 5 repeated doses of 20 µM, or a cumulative single dose of 100 µM. Data are shown as individual data points, mean, and SD. N = 4 hydrogels per condition. *p<0.5, **p<0.01, ***p<0.001.

8MGBA: For 8WT GBM cells treated with a single dose of 5 µM, 5 repeated doses of 5 µM, or a cumulative single dose of 25 µM, we observed a decrease in the metabolic activity with dose (Fig. 2B). The 25 µM treatment was most effective in decreasing metabolic activity of 8WT with this treatment being significantly more effective than DMSO vehicle control as well as more effective than 5 µM treatment (Fig. 2B). However, both treatments of 5 µM and 5 repeated doses of 5 µM were not statistically significant from the DMSO control group, with 5x5 µM group being marginally significant compared to DMSO control, p-value = 0.08 (Fig. 2B). When 8TMZres were treated with either a single dose of 5 µM, 5 repeated doses of 5 µM, or a cumulative single dose of 25 µM, we did not observe a statistically significant decrease in the metabolic activity (Fig. 2B). Interestingly, the metabolic activity of the 5x5 µM treated 8TMZres cell were significantly higher compared to the group treated with a single dose of 5 µM (Fig. 2B). Next, when 8WT were treated with either a single dose of 20 µM, 5 repeated doses of 20 µM, or a cumulative single dose of 100 µM, these three treatments all significantly decreased cell metabolic activity (Fig. 2C). However, we did not observe any significant differences in response between these higher TMZ dose treatments (Fig. 2C). When 8TMZres were treated with either a single dose of 20 µM, 5 repeated doses of 20 µM, or a cumulative single dose of 100 µM, we observed that the 100 µM TMZ treatment decreased the metabolic activity of 8TMZres cells compared to DMSO control (Fig. 2C) and the 5 x 20 µM metronomic treatment (Fig. 2C).

42MGBA: We observed a significant decreased in the metabolic activity of 42WT cells treated with a single dose of 5 µM, 5 repeated doses of 5 µM, or a cumulative single dose of 25 µM (Fig. 2D). However, these low TMZ dose treatments were not statistically different from each (Fig. 2D). When 42TMZres were treated with either a single dose of 5 µM, 5 repeated doses of 5 µM, or a cumulative single dose of 25 µM, we did not observe a statistically significant decrease in the metabolic activity of 42TMZres (Fig. 2D). The metabolic activity of the 5x5 µM treated 42TMZres cell were significantly higher compared to the group treated with a single dose of 25 µM (Fig. 2D). Similarly, 42WT treated with either a single dose of 20 µM, 5 repeated doses of 20 µM, or a cumulative single dose of 100 µM all displayed significantly decreased metabolic activity (Fig. 2E). Interestingly, 5x20 µM treatment was most effective in decreasing the metabolic activity of 42WT (Fig. 2E). Interestingly, 42TMZres cells did not display a significant difference in response between dose groups (single dose of 20 µM, 5 repeated doses of 20 µM, or a cumulative single dose of 100 µM) or compared to the DMSO control (Fig. 2E)

### Acquired Temozolomide Resistance Alters Apoptosis Pathways

We analyzed a panel of apoptosis-related proteins in temozolomide (TMZ)-sensitive vs. TMZ-resistant glioblastoma (GBM) cell lines after treatment with DMSO vehicle control or temozolomide (TMZ) at the appropriate GR50 concentration (as determined for 7 days culture, Table 1) for each cell line. We profiled the relative expression of 35 apoptosis-related proteins and observed that TMZ treatment significantly affected expression of most of the profiled apoptosis-related proteins (Fig 3A, Fig 4A, Fig S5). Notably, all four cell lines, regardless of treatment and TMZ sensitivity status, expressed significantly high levels of Pro-Caspase-3 (Pro-CASP3/PC-3) (Fig. 3B and Fig. 4B). GR50 TMZ treatment resulted in decreased expression of Pro-CASP3 in 8MGBA pair and increased expression of Cleaved CASP3 (Fig. 3B). However, in 42MGBA pair, GR50 TMZ treatment resulted in increased expression of Pro-CASP3 (with higher expression in 42TMZres), while expression of Cleaved-CASP3 followed similar trend to 8MGBA pair (Fig. 4B). Moreover, we observed high expression of phosphorylated p53 (Phospho-p53) S15, S46, and S392 (Fig. 3C and Fig. 4C). Most strikingly, we saw highly significant increase in expression of Phospho-p53, S46 in 42MGBA-TMZres vs. 42MGBA-WT with GR50 TMZ treatment further increasing Phospho-p53, S46 expression in 42MGBA-TMZres (Fig. 4C). We observed a similar trend in the expression of Phospho-p53 S392 in 42MGBA pair (Fig. 4C). Collectively, we observed TMZ resistance leads to altered apoptosis pathways, specifically shifts in the expression of multiple apoptosis-related proteins in the TMZ-treated WT vs. TMZres cells.

**Fig. 3.**
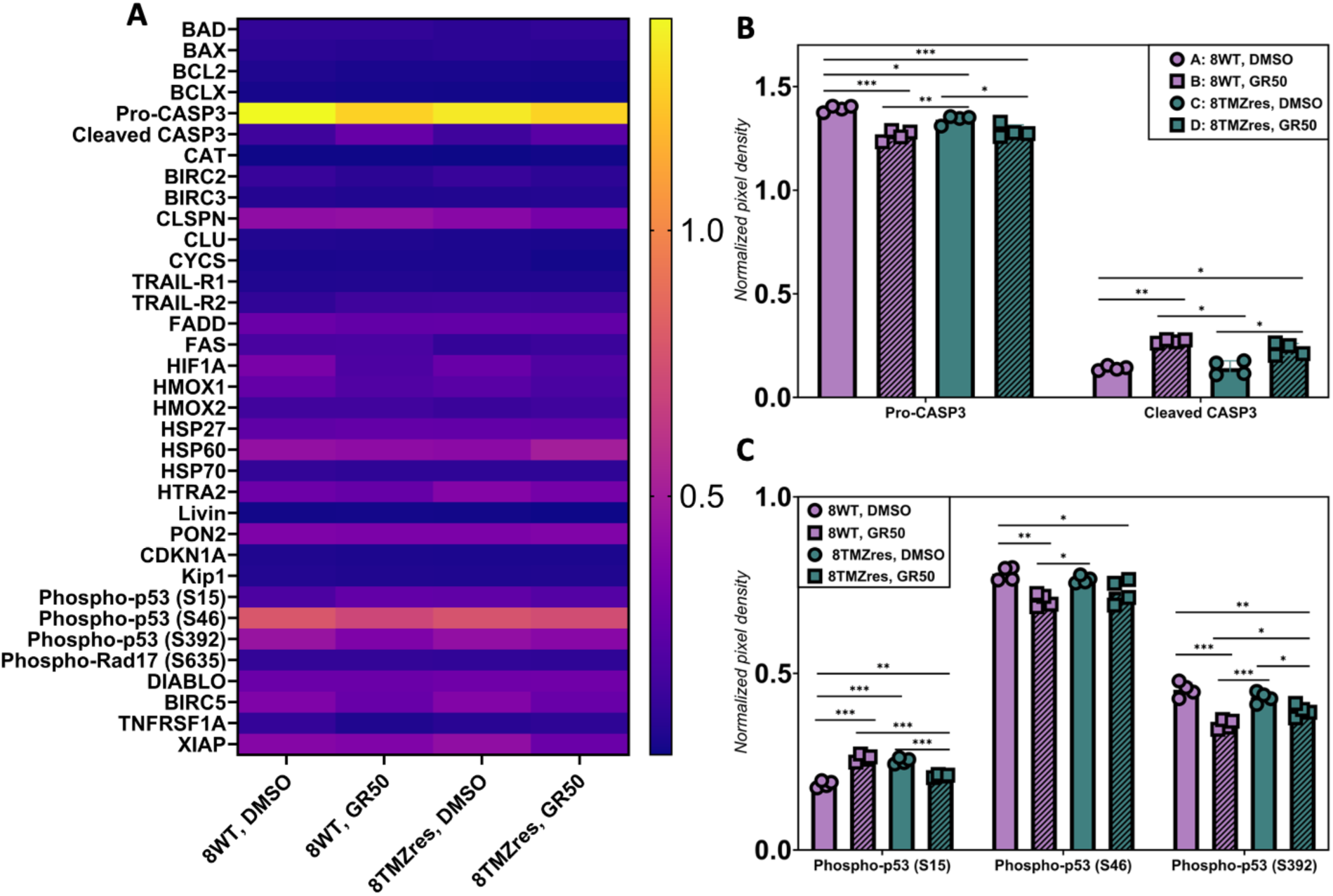
Expression of apoptosis-related proteins in the 8MGBA pair. **(A)** Heat map showing the normalized relative expression of 35 apoptosis-related proteins in 8MGBA-WT vs. TMZ-res cells after treatment with either DMSO vehicle control or TMZ at GR50, D7 doses for 7 days. **(B)** Relative expression levels of Pro-Caspase-3 (Pro-CASP3/PC-3) and cleaved CASP3. **(C)** Relative expression levels of Phosopho-p53 (S15), Phosopho-p53 (S46), Phosopho-p53 (S392). Data are shown as individual data points, mean, and SD. N = 4 hydrogels per condition. *p<0.5, **p<0.01, ***p<0.001.

**Fig. 4.**
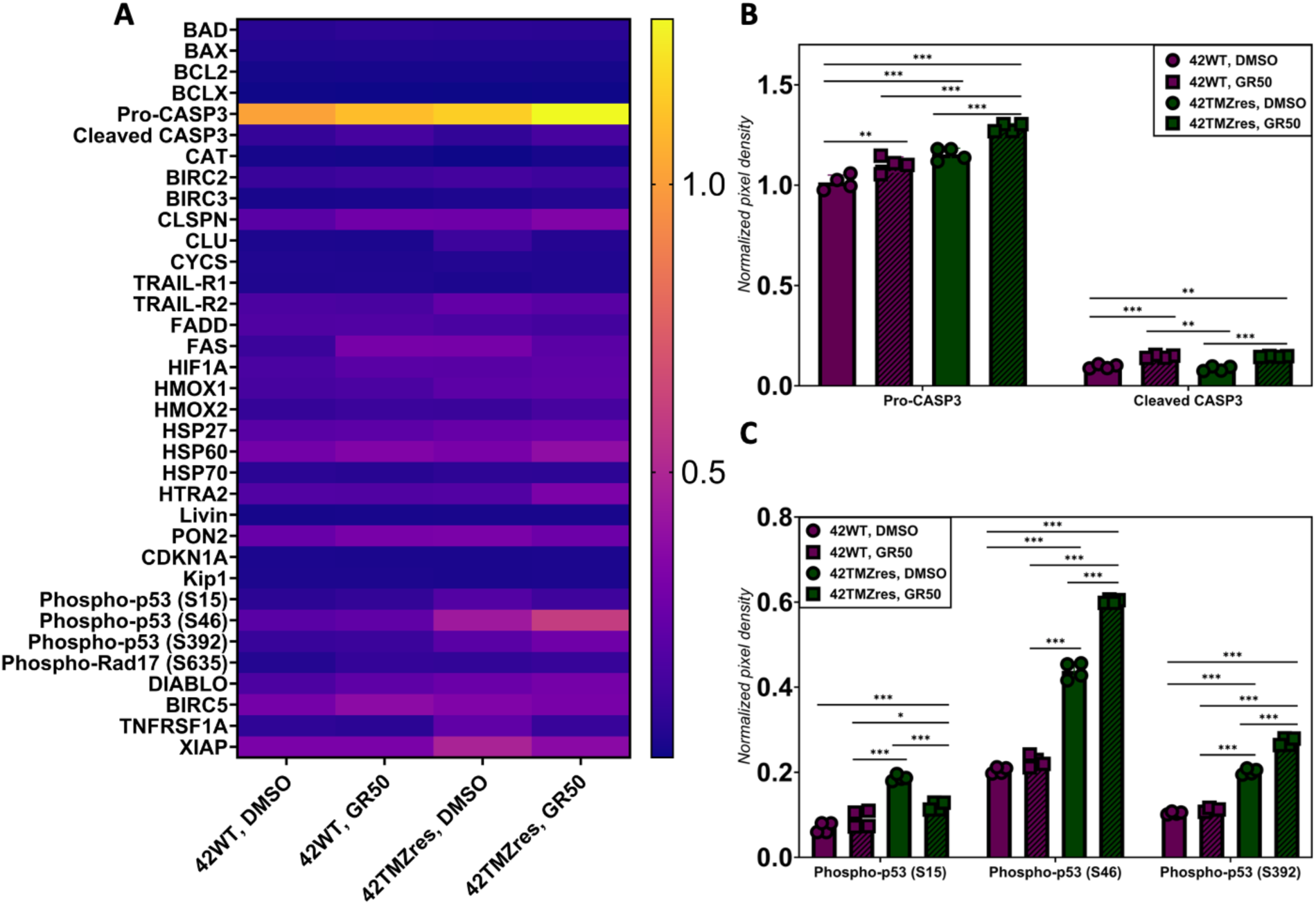
Expression of apoptosis-related proteins in the 42MGBA pair. **(A)** Heat map showing the normalized relative expression of 35 apoptosis-related proteins in 42MGBA-WT vs. TMZ-res cells after treatment with either DMSO vehicle control or TMZ at GR50, D7 doses for 7 days. **(B)** Relative expression levels of Pro-Caspase-3 (Pro-CASP3/PC-3) and cleaved CASP3. **(C)** Relative expression levels of Phosopho-p53 (S15), Phosopho-p53 (S46), Phosopho-p53 (S392). Data are shown as individual data points, mean, and SD. N = 4 hydrogels per condition. *p<0.5, **p<0.01, ***p<0.001.

### Acquired Temozolomide Resistance Leads to Reduced Motility in A Three-Dimensional Spheroid Invasion Model

The invasive capacity of temozolomide (TMZ) resistant vs. responsive 8MGBA and 42MGBA glioblastoma (GBM) cell pairs was compared using a cell spheroid-based invasion assay without presence of TMZ in culture. Radial patterns of cell motility for cells migrating into the gelatin hydrogel from cell spheroids generated from 5000 GBM cells were evaluated for up to 3 days (Fig. 5B). Notably, both TMZ resistant variants, 8MGBA-TMZres and 42MGBA-TMZres, displayed significantly reduced spheroid outgrowth areas compared to their parental, TMZ-sensitive controls (Fig. 5C and Fig 5D). The most TMZ resistant cell line, 42MGBA-TMZres, showed minimal invasion compared to all three other cell lines (Fig. 5D).

**Fig. 5.**
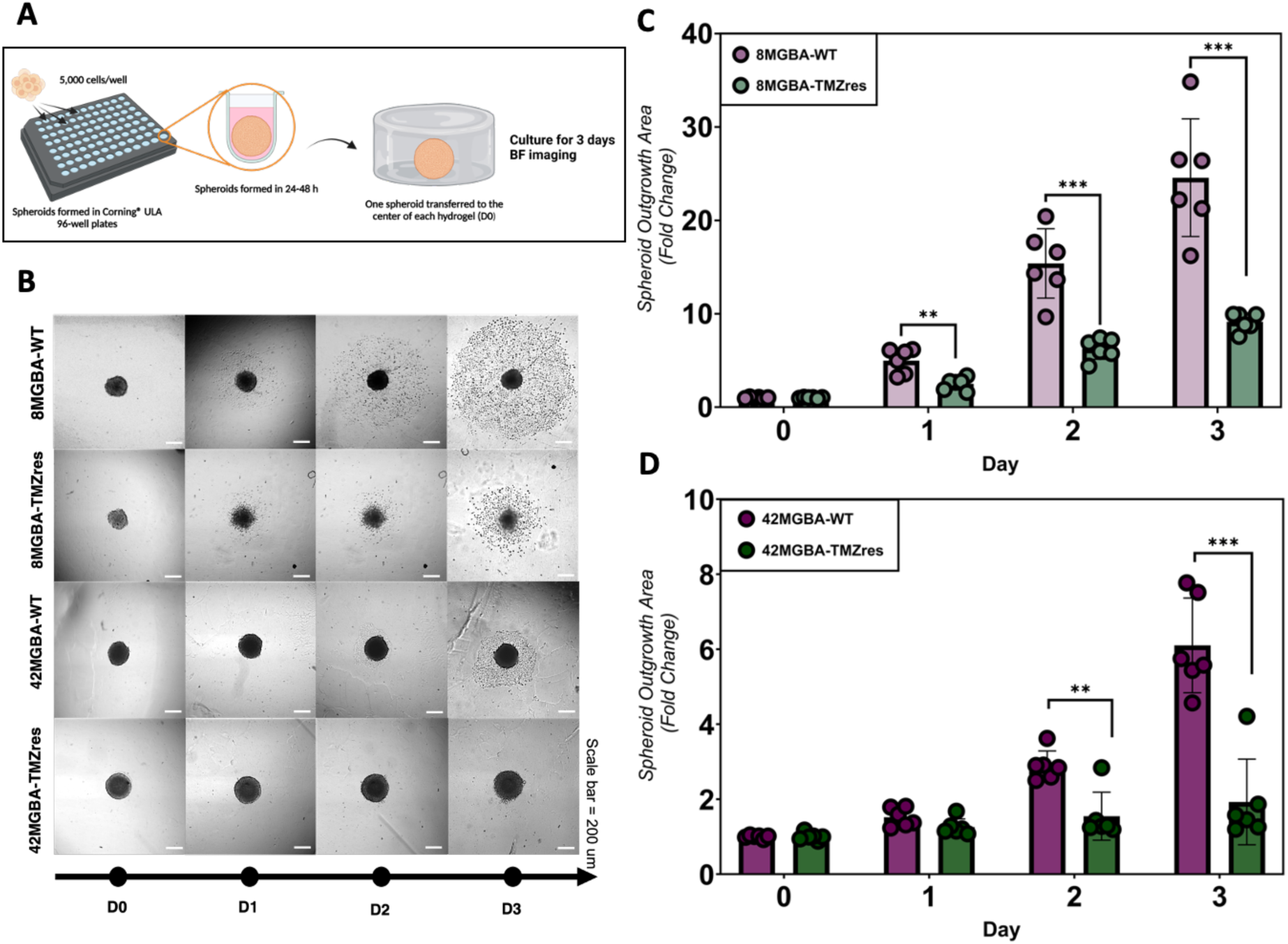
Spheroid invasion model to quantify the effects of acquired TMZ resistance on glioblastoma cell motility. **(A)** Experimental setup to determine cell motility in TMZ-sensitive vs. resistant GBM cell lines. Each spheroid was formed by culturing 5000 GBM cells in ultra-low attachment plates; individual spheroids were then encapsulated in GelMA hydrogels. Radial patterns of cell motility of cells migrating into the hydrogel from spheroids were evaluated for 3 days by taking bright-field (BF) images. **(B)** Representative BF images of cell spheroid outgrowth. **(C)** Fold change of spheroid outgrowth area of 8MGBA-WT vs. 8MGBA-TMZres cell lines. **(D)** Fold change of spheroid outgrowth area of 8MGBA-WT vs. 42MGBA-TMZres cell lines. Data are shown as individual data points, mean, and SD. N = 6 hydrogels per condition. **p<0.01, ***p<0.001.

### Temozolomide-Resistant GBM Cells Display Differential Expression of Pro-Angiogenic and Matrix-Remodeling Associated Cytokines

Finally, we examined a panel of 55 angiogenesis-related soluble factors in the secretome of the temozolomide (TMZ)-sensitive vs. TMZ-resistant glioblastoma (GBM) cells using a cytokine array. Invading GBM cells induce significant remodeling events; TMZ-sensitive vs. TMZ-resistant GBM cells secreted a variety of remodeling and angiogenesis-related proteins over the course of 7 days cultured in vitro in GelMA hydrogels (Fig. S8, Supporting Information). We quantified mean pixel density of cytokine spots normalized to the positive control (Fig. S8, Supporting Information). Many profiled cytokines had negligible expression (comparable to the negative control and less than 5% of positive control); therefore, we chose to further analyze cytokines that were detected robustly. We observed significant expression of 15 cytokines in the 8MGBA pair (Fig. 6A and Fig. S9) and 17 in the 42MGBA pair (Fig. 7A and Fig. S9, Supporting Information). Out of the most detectable cytokines, we saw significant differences in expression between TMZ-sensitive vs. TMZ-resistant groups, of the following soluble factors: (a) 8MGBA-WT vs. 8MGBA-TMZres, 4 cytokines: IGFBP-3, IL-8, Serpin F1, and VEGF and (b) 42MGBA-WT vs. 42MGBA-TMZres, 11 cytokines: ANGPT1, CXCL16, Endostatin (Collagen XVIII), IGFBP-2, IL-8, MCP-1, PTX3, PDGFA, Serpin E1, Serpin F1, uPA (PLAU) (Fig. 6A, Fig. 7A, Fig. S9, Supporting Information).

**Fig. 6.**
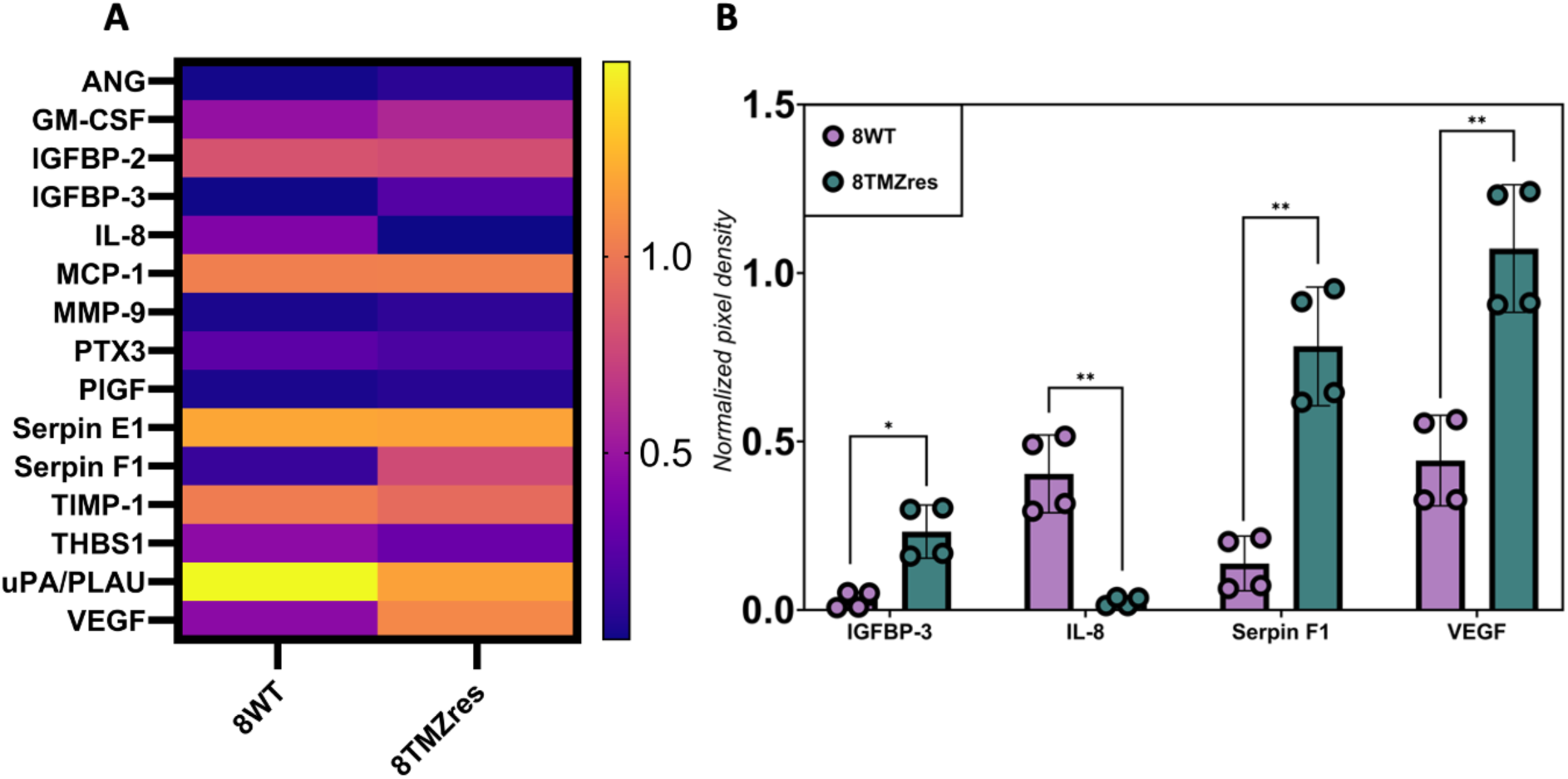
Expression of matrix remodeling and angiogenesis-related cytokines in the 8MGBA pair. **(A)** Heat map showing the normalized relative expression of 15 significantly detected matrix remodeling and angiogenesis-related cytokines in the secretome collected from 8MGBA-WT vs. TMZ-res cells after 7 days of culture in GelMA hydrogels. **(B)** Relative expression levels of IGFBP-3, IL-8, Serpin F1, and VEGF. Data are shown as individual data points, mean, and SD. N = 4 hydrogels per condition. *p<0.5, **p<0.01, ***p<0.001.

**Fig. 7.**
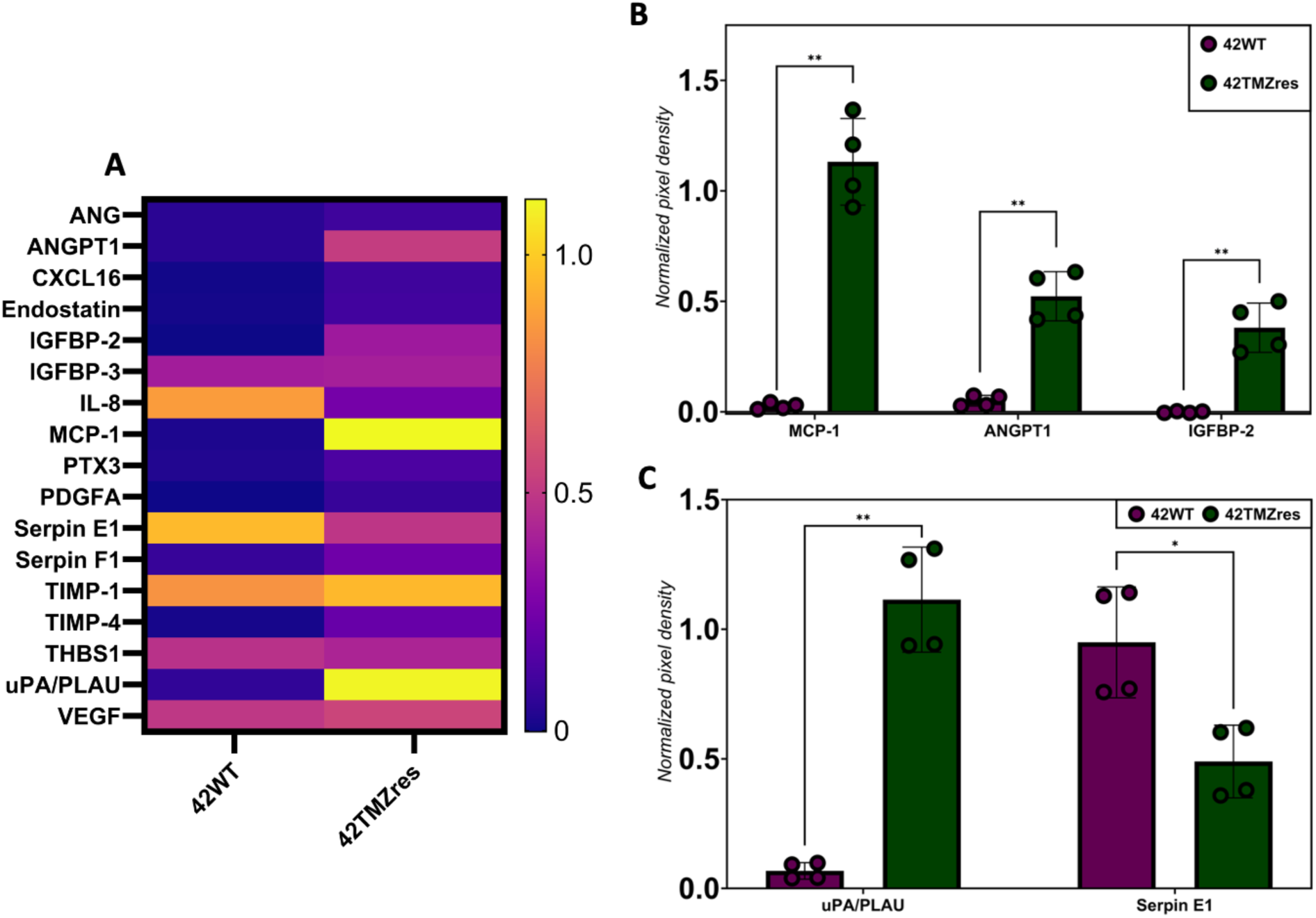
Expression of matrix remodeling and angiogenesis-related cytokines in the 42MGBA pair. **(A)** Heat map showing the normalized relative expression of 17 significantly detected matrix remodeling and angiogenesis-related cytokines in the secretome collected from 42MGBA-WT vs. TMZ-res cells after 7 days of culture in GelMA hydrogels. **(B)** Relative expression levels of MCP-1, ANGPT-1, IGFBP-2. **(C)** Relative expression levels of uPA/PLAU and Serpin E1. Data are shown as individual data points, mean, and SD. N = 4 hydrogels per condition. *p<0.5, **p<0.01, ***p<0.001.

Acquired TMZ resistance resulted in significant upregulation of IGFBP-3 Serpin F1, and VEGF and significant downregulation of IL-8 in the 8MGBA pair (Fig. 6B). Notably, both 8MGBA-WT and 8MGBA-TMZres secreted high levels of GM-CSF, IGFBP-2, MCP-1, Serpin E1, TIMP-1, THBS1, and uPA, although differences in expression of these cytokines were not statistically significant between 8WT and 8TMZres (Fig. 6A and Fig. S9, Supporting Information).

Interestingly, the changes in 42MGBA-TMZres vs. 42MGBA were different comparing to the 8MGBA pair; in this pair, we observed upregulation of different pro-angiogenic factors with acquired TMZ resistance: ANGPT1, CXCL16, Endostatin, IGFBP-2, MCP-1, PTX3, PDGFA, Serpin F1, uPA (Fig. 7A, Fig. 7B, and Fig. S9, Supporting Information). Acquired TMZ resistance resulted in significant downregulation of IL-8 (Fig. S10, Supporting Information, consistent with 8MGBA) and Serpin E1 (Fig. 7B). Notably, both 42MGBA-WT and 42MGBA-TMZres secreted high levels of IGFBP-2, TIMP-1, THBS1, and VEGF, although differences in expression of these cytokines were not statistically significant between 42WT and 42TMZres (Fig. 7A and Fig. S9, Supporting Information).

## Discussion

The current glioblastoma (GBM) treatment strategy provides minimal benefit to patients, even when treated with highly aggressive chemotherapy. The standard-of-care alkylating agent, temozolomide (TMZ), ultimately fails to kill residual and highly invasive tumor cells after surgical resection and radiotherapy. Even when patients initially show a positive response to TMZ therapy, the tumor rapidly acquires resistance to TMZ during treatment. As the result, TMZ therapy only adds about 4 months to the median overall survival of GBM patients (4, 8, 33). Due to the absence of FDA-approved second-line treatments for GBM, it is crucial to develop more clinically relevant in vitro models of TMZ-resistant GBM. Currently, most in vitro models to evaluate GBM drug response fail to consider the effects of the extracellular matrix and tumor microenvironment (19, 34–36). Importantly, the effects of acquired temozolomide resistance on GBM behavior in three-dimensional (3D) in vitro models of the brain microenvironment are yet to be defined. Understanding how the standard-of-care drug treatment shapes the phenotypic and molecular changes of GBM progression within the 3D extracellular matrix models will reveal key insights into discovery of new therapeutic targets.

In this study, we developed a robust 3D in vitro model of acquired TMZ resistance using a set of isogenically-matched TMZ-sensitive vs. TMZ-resistant GBM cell lines in gelatin hydrogels. Gelatin presents an attractive platform for engineering the extracellular matrix environment. Being a natural polymer derived from collagen, gelatin contains cell adhesion and degradation sites which facilitate cell-initiated matrix remodeling events (37). While collagen is not abundant in healthy brain tissue, it has been shown to be significantly present in GBM (38). Through the addition of methacrylate groups to the gelatin’s amine-containing side groups, gelatin can be functionalized into gelatin methacryloyl (GelMA), which exhibits improved mechanical properties that can be tuned to mimic the biophysical and biochemical properties of the brain tissue (39). Moreover, the macroscale of our GelMA platform enables the collection of larger sample volumes for subsequent high-throughput -omics analyses. Importantly, we have previously developed GelMA-based platforms that contain engineered brain microvasculature and the perivascular niche (PVN) as well as gradients of tumor-relevant hyaluronic acid (11, 13, 23, 25–27, 39). Therefore, future studies can leverage the potential to incorporate additional extracellular matrix proteins (e.g., hyaluronic acid) and to include PVN microenvironments to investigate synergistic effects of the perivascular and extracellular matrix-derived signals on drug response.

Here, we first demonstrate that our GelMA-based engineered extracellular matrix model supports robust detection of drug response between drug-sensitive and drug-resistant GBM cells. We defined detection limits of TMZ response using two isogenically-matched TMZ-responsive (WT) and TMZ-resistant (TMZres) 8MGBA and 42MGBA cell lines. We showed drastic changes in relative growth rates for up to 7 days post single-dose TMZ treatment. In fact, our data suggest that it is beneficial to monitor TMZ response for 7 days post treatment, a potential advantage for 3D culture over 2D systems where cell confluence can be a confounding factor. Indeed, we show that 8MGBA-TMZres start exhibiting their resistant behavior 5 days after treatment, while 42MGBA-TMZres show resistance as early as 3 days post treatment. The 42MGBA pair is particularly interesting.

The difference in GR50 values 7 days post treatment is >1000 µM for 42MGBA-WT vs. 42MGBA-TMZres. In fact, 42MGBA-WT is quite sensitive to TMZ treatment with GR50 values <10 µM measured at days 3, 5, and 7, while GR50 values of the 42MGBA-TMZres reach >1000 µM. Interestingly, even when treated with very high TMZ doses (>300 µM), 42MGBA cells do not display any true cytotoxic effects, i.e., their relative growth rates do not go below zero. These results are consistent with previous findings by Tiek et al., which showed that TMZ did not induce PARP cleavage (apoptotic marker) in 42MGBA-WT cells, while PARP cleavage was robustly induced by in 8MGBA-WT (28). It is important to note that even 10 µM would be considered a very high concentration for any new generation drug; however, TMZ is administered to patients using an aggressive treatment strategy (40). While patient undergo radiation therapy, they are concurrently given daily oral TMZ at 75 mg/m2 body surface area (BSA)/day for 6 weeks, then followed by adjuvant TMZ treatment with 6 maintenance cycles at 150–200 mg/m2 BSA/day for the first 5 days of a 28-day cycle (3, 4). There have been multiple studies with the goal to estimate clinically relevant TMZ concentrations in vivo (peak concentrations in the brain interstitium/CSF), and numbers ranging between 0.82 µM to 34.54 µM have been suggested, with around 5 µM appearing to be the clinically relevant dose. However, most in vitro studies use high TMZ concentrations in the range of 100 µM and as high as 4 mM (41, 42). As a result, there is a limited number of in vitro studies that evaluate the effect of low physiologically relevant TMZ concentrations.

We also explored the feasibility of examining cell response to frequent and repetitive administration of low dose (metronomic) TMZ, which may be particularly useful using a 3D model. While prior studies have suggested that single-dose TMZ treatments at concentrations around 5 µM have minimal effect on apoptosis of GBM cell lines, we sought to evaluate the effect of low-concertation single and repeated TMZ doses using our WT vs. TMZres cells and gelatin hydrogel model (43, 44). We chose to treat our isogenically-matched GBM cells with the following TMZ dosing schedules: (1) a single dose of 5 µM, or 5 repeated doses of 5 µM, or a cumulative single dose of 25 µM; or (2) a single dose of 20 µM, or 5 repeated doses of 20 µM, or a cumulative single dose of 100 µM. We selected 5 µM (optimal dose) and 20 µM (slightly high dose) doses to validate our model at physiologically relevant TMZ concentrations. We were further interested to see how these single low doses would perform compared to 5 daily doses of 5 µM and 20 µM. Additionally, we included single high doses of TMZ at 25 µM and 100 µM to represent single cumulative doses of the 5 low-dose treatments. Previous studies have shown that repeated low doses of TMZ induce similar cytotoxic effects compared to a single high dose (44). Our data support a similar trend; however, due to the nature of our cell models (TMZ-sensitive vs acquired resistance), we observe significant variability between cell lines. Additionally, we chose to measure metabolic activity of the cells, which is an inherently surrogate measure of drug cytotoxicity (45, 46). As expected, these low dose treatments were largely ineffective to induce any cytotoxicity in the TMZ-resistant cell lines; however, these treatments were effective in inducing significant decrease in metabolic activity of the TMZ-sensitive cell lines. Collectively, these results show that our model can be successfully used to study the effects of physiologically relevant drug treatments.

We were then interested to see how single-dose TMZ treatment affected expression of apoptosis-related proteins in our model. We chose to treat 8MGBA-WT/TMZres and 42MGBA-WT/TMZres with TMZ concentrations at the GR50 values (day 7) that we previously determined. These concentrations were: 28 µM for 8MGBA-WT, 279 µM for 8MGBA-TMZres, 3 µM for 42MGBA-WT, 1225 µM for 42MGBA-TMZres. In this experiment, we demonstrate a path to effectively compare drug-induced molecular changes in 3D-cultured GBM cells. Specifically, by first using hydrogel models to identify GR50 values from cytotoxic effects of TMZ treatment, we examine differences in TMZ-induced molecular changes in the GBM cells for each line’s unique GR50 value. Our data showed that all four cell lines, regardless of TMZ treatment, expressed high levels of Pro-Caspase-3 (Pro-CASP3/PC-3). In fact, PC-3 expression was generally above the assay’s positive control. PC-3, the precursor/immature zymogen to Caspase-3 (CASP3), is a pro-apoptotic protein. Generally, apoptosis is believed to take place when PC-3 is cleaved to CASP3 (47, 48). In our study, we observed increase in the relative expression of cleaved CASP3 as the result of TMZ treatment at the GR50 concentrations for all four profiled cell lines. This result suggests that the GR50 treatments indeed initiated cellular apoptosis, further validating our dose-response studies. It is important to note that PC-3 expression was significantly higher than cleaved CASP3 expression. Particularly in the 42MGBA pair, we observed increase in PC-3 expression with TMZ treatment with 42MGBA-TMZres, GR50-treated cells having the highest expression of PC-3. Recently, there has been an increasing number of studies suggesting that the PC-3 is overexpressed in many tumors compared to the matched normal tissues (49–51). Importantly, PC-3 has been shown to be abnormally high in all brain tumors, including GBM (52–54). The overexpression of this pro-apoptotic marker in cancer is rather paradoxical; however, recent studies suggest that PC-3 activity may have non-apoptotic roles and be involved in oncogenic transformations (55). In fact, PC-3 is an emerging therapeutic target, with the PC-3 targeting compound, PAC-1, currently being investigated in clinical trials in patients with end-stage cancers (56).

We observed high expression of phosphorylated p53 at serines 15, 45, and 392 (Phospho-p53 S15/S46/S392) in response to GBM cell exposure to GR50 dose levels. Known as the “guardian of the genome,” the tumor suppressor p53 has been shown to activate numerous cellular programs in cancer such as cell cycle arrest, apoptosis, differentiation, DNA repair, autophagy, and senescence through complex signaling pathways (57). However, despite numerous studies, the prognostic value of p53 in GBM remains under debate (58). Similarly, there is no consensus on the relationship between p53 and TMZ (59). Among multiple mechanisms controlling p53 function, post-translational modifications (PTMs), such as phosphorylation, are key (60). In our study, we did not observe any clear patterns in Phospho-p53 expression in the 8MGBA isogenic pair; however, we observed significant changes in Phospho-p53 expression in the 42MGBA isogenic pair. Specifically, 42MGBA-TMZres expressed significantly higher levels of Phospho-p53 S15, S46, S392 in response to TMZ treatment at the GR50 dose. TMZ treatment of 42MGBA-TMZres decreased Phospho-p53 S15 expression and significantly increased Phsopho-p53 at both S46 and S392, though Phospho-p53 expression in 42MGBA-WT was unaffected by TMZ treatment. Phosphorylation of p53 at serine 15 (S15) is believed to lead to cell cycle arrest; whereas phosphorylation of p53 at S46 is a crucial occurrence that drives cells to choose apoptosis over cell cycle arrest; phosphorylation of p53 at S392 stabilizes p53 tetramers and boosts DNA binding activity (61, 62). Both 8MGBA and 42MGBA harbor TP53 mutations. The homozygous R273C in 8MGBA is a well-established hotspot mutation that impairs p53 DNA binding, while the heterozygous R282Q in 42MGBA is a less-common amino acid change at a residue that, when mutated, can change p53 conformational structure (63–66). Interestingly, experimental studies have shown that R282Q retains p53 wild type activities, which may partly explain why TMZ induces p53 phosphorylation events associated with cell cycle arrest and apoptosis selectively in the 42MGBA cell models (67). While the different tenor in responses between TMZ resistant and WT GBM cells is also likely due to their significantly different GR50 dose value (28 µM for 8MGBA-WT, 279 µM for 8MGBA-TMZres, 3 µM for 42MGBA-WT, 1225 µM for 42MGBA-TMZres; overall, we show that our hydrogel model is suitable to identify then study profile molecular changes in drug-treated GBM in response to a wide range of TMZ doses.

Having demonstrated robust alterations in drug response within our model of acquired TMZ resistance, we wished to further demonstrate phenotypical shifts in GBM behavior resulting from acquired TMZ resistance. Previously, Tiek et al. used a two-dimensional scratch assay to show 8MGBA-TMZres cells were significantly more migratory than 8MGBA-WT cells, while 42MGBA-TMZres showed a modest but nonsignificant reduction in cell migration compared to 42MGBA-WT (28). We performed a three-dimensional spheroid invasion assay using the 8MGBA and 42MGBA isogenically matched pairs in control media. Interestingly, our results suggest that 8MGBA-WT are significantly more migratory than 8MGBA-TMZres. Similarly, our data show that 42MGBA-WT are more migratory than 42MGBA-TMZres. Thus, our data suggest that acquired drug resistance leads to decrease in the migratory potential of GBM cells within GelMA-based extracellular matrix hydrogels. GelMA contains MMP-degradable sites that aid in cellular migration through matrix degradation. Previous research conducted in our lab has confirmed the expression of MMP2 and MMP9 by GBM cells when cultured within GelMA hydrogels, suggesting opportunities to understand whether the observed decrease in cell motility may be tied to molecular changes in matrix remodeling capacity (13, 27, 29). It is important to note that we monitored cellular migration from spheroids for only 3 days and in the absence of TMZ; therefore, ongoing studies are focusing on establishing a long-term spheroid assay to further investigate TMZ-resistant GBM invasion and matrix remodeling in response to both physiological and GR50 level TMZ doses.

Finally, we assessed a panel of soluble factors related to extracellular matrix remodeling and angiogenesis. We profiled secretome of 8MGBA-WT/TMZres and 42MGBA-WT/TMZres cells in the absence of TMZ treatment against a panel of 55 angiogenesis-related cytokines. The invasive GBM cells induce significant matrix remodeling events; therefore, we were interested to evaluate the effect of acquired TMZ resistance on the expression of matrix remodeling-associated cytokines (11). We observed significant expression of 15 cytokines in the 8MGBA pair and 17 in the 42MGBA pair. While some matrix remodeling cytokines were highly expressed regardless of TMZ sensitivity status, several cytokines were either significantly upregulated or downregulated as the result of acquired TMZ resistance. Interestingly, we observed significantly increased expression of the Urokinase Plasminogen Activator (uPA) and significantly decreased expression of the Plasminogen Activator Inhibitor-1, known as Endothelial Plasminogen Activator Inhibitor (SERPINE1), in 42MGBA-TMZres compared to 42MGBA-WT. The expression of uPA and SERPINE1 was very high in both 8MGBA-WT and 8MGBA-TMZres. There has been increasing evidence supporting the role of the plasminogen activator system, i.e., uPA (and its receptor uPAR) and SERPINE1, in multiple cancers. Overexpression of uPA and SERPINE1 enhances tumor cell migration and invasion and is correlated with poor prognosis. Both uPA and SERPINE1 are directly associated with triggering the epithelial-to-mesenchymal transition as well as the acquisition of stem-cell like characteristics, and the development of therapeutic resistance (68, 69).

Analysis of the secreted factors also revealed significant increase in the expression of MCP-1, ANGPT1, and IGFBP-2 in 42MGBA-TMZres compared to 42MGBA-WT. MCP-1 and IGFBP-2 expression was high in both 8MGBA-WT and 8MGBA-TMZres. Monocyte chemotactic protein-1 (MCP-1) is associated with tumor development, tumor invasion and metastasis, angiogenesis, and immune cell infiltration; cancer-secreted MCP-1 has been correlated with a poor clinical outcome, due to the induction of tumor-associated macrophage infiltration and tumor invasion/metastasis in several solid tumors (70). Insulin-like growth factor binding protein 2 (IGFBP-2) has been shown to promote GBM cell migration and invasion and to contribute to cancer progression, recurrence, and poor survival (71). Moreover, our data show significant secretion of other matrix remodeling cytokines by GBM cells, namely, VEGF, TIMP-1, and THBS1. Vascular endothelial growth factor (VEGF) is the key mediator of angiogenesis in GBM (72). Our data show that 8MGBA-TMZres secrete significantly higher levels of VEGF compared to their parental 8MGBA-WT, while both 42MGBA-WT/TMZres secrete much VEGF. Tissue inhibitors of metalloproteinase (TIMP) family proteins are peptidases involved in extracellular matrix degradation. TIMPs can indirectly regulate remodeling of the ECM by regulating matrix metalloproteinase (MMP) activity (73). Matricellular protein Thrombospondin-1 (THBS1) may promote angiogenesis and tumor cell invasion (74). We additionally observed increased secretion of ANGPT1 in 42MGBA-TMZres compared to 42MGBA-WT. Angiopoietin 1 (ANGPT1) may modulate cell-cell and cell-matrix associations and may promote the differentiation phase of angiogenesis (75). Thus far, the discussed cytokines associated with increased cancer cell invasion were mostly upregulated in the expression of TMZ-resistant cell lines within our model. Yet, the results of our spheroid invasion model suggest that acquired TMZ resistance leads to decreased migration. Interestingly, IL-8 expression was downregulated in both 8MGBA-TMZres and 42MGBA-TMZres compared to their parental WT cell lines. Interleukin-8 (IL-8) has garnered considerable interest as a factor that promotes migration and angiogenesis in various cancers, and it might also play a role in regulating functions of GBM cancer stem cells (CSCs). Decreased secretion of IL-8 by TMZ-resistant cells might be one possible explanation of our rather counter-intuitive results, where TMZ-resistant cells show decreased migration compared to their parental wild-type cells. Hence, our ongoing research is dedicated to gaining a deeper understanding of matrix remodeling and the migratory capabilities of TMZ-resistant GBM cells.

This endeavor seeks to establish connections between observed therapeutic behaviors and clinical outcomes, ultimately contributing to the development of more effective treatments for GBM patients. Using a robust engineered extracellular-matrix model, we have characterized the behavior of four glioblastoma (GBM) cell lines with well-defined shifts in response to the front-line chemotherapeutic, temozolomide (TMZ). While some aspects of the observed effects of acquired TMZ resistance on GBM cell behavior agree with several in vitro and in vivo studies, we acknowledge that observations made with these two isogenically-matched cell lines cannot be extrapolated to all GBM tumor cells due to the inherent heterogeneity that exists between patients. However, our platform allows the investigation of tumor cell behavior in a patient-specific manner, thereby enabling the development and screening of personalized therapies. Future work in our laboratory will focus on profiling a larger panel of cell lines and patient-derived xenograft lines within our glioblastoma tumor microenvironment models to correlate observed therapeutic phenotypes to known clinical outcomes.

## Materials and Methods

### Cell Culture

8MGBA-WT, 8MGBA-TMZres, 42MGBA-WT, and 42MGBA-TMZres cells were provided by Dr. Rebecca B. Riggins (Lombardi Comprehensive Cancer Center (LCCC), Georgetown University, Washington DC). All cells tested negative for Mycoplasma contamination using MycoStrip™ – Mycoplasma Detection Kit (InvivoGen, San Diego, CA). Cells were grown in adherent culture flasks in DMEM (Cat. #11965, Thermo Fisher Scientific, Waltham, MA) supplemented with 10% FBS (R&D Systems, Minneapolis, MN) and 0.2% v/v plasmocin (InvivoGen, San Diego, CA). Growth media used to culture 8MBGA-TMZres was further supplemented with 100 µM temozolomide (TMZ) to maintain TMZ resistance in 8MBGA-TMZres. TMZ (Cat. #S1237, Selleckchem, Houston, TX) was dissolved in DMSO (Thermo Fisher Scientific, Waltham, MA) to a stock concentration of 130 mM and subsequently added to 8MGBA-TMZres growth media at the indicated concentration shortly prior to adding media to cells. All cells were passaged fewer than 10 times and maintained in a humidified incubator with 95% air and 5% CO2 at 37 °C.

### Methacrylamide-Functionalized Gelatin (GelMA) Synthesis and Characterization

GelMA was synthesized as previously described (29, 30). Briefly, 1 g of porcine gelatin type A, 300 bloom (Sigma Aldrich, St. Louis, MO) was dissolved in 10 mL of carbonate-bicarbonate (CB) buffer (pH 9.4) at 50 °C. Subsequently, 40 µL of methacrylic anhydride (Sigma Aldrich, St. Louis, MO) was added dropwise, and the reaction proceeded for 1 hour with vigorous stirring (400 RPM). The reaction was quenched with 40 mL of warm deionized water and dialyzed in 12-14 kDa dialysis membranes for 7 days against deionized water with daily water exchange. The product was then frozen and lyophilized. 1HNMR was used to determine the degree of functionalization (DOF). GelMA with DOF of ∼55% was used in this study. Compressive moduli of GelMA hydrogels were measured using an Instron 5943 mechanical tester (Norwood, MA). Hydrogels were tested under unconfined compression at the rate of 0.1 mm/min, with the Young’s modulus obtained from the linear region of the stress-strain curve (2.5–17.5% strain) using a custom MATLAB (MathWorks, Natick, MA) code.

### Fabrication of Cell Laden GelMA Hydrogels

Glioblastoma cells of interest (8MGBA, 8MGBA-TMZres, 42MGBA, or 42MGBA-TMZres) were homogeneously resuspended in the GelMA precursor solution at the concentration of 5 x 105 cells/mL, and the resulting cell suspension was pipetted into custom Teflon molds (5 mm diameter, 1 mm thick). Hydrogels were formed after photopolymerization for 45 seconds using a UV lamp (λ = 365 nm, 5.69 mW/cm2). Hydrogels were deposited into 48-well plates with each well containing 500 µL of growth media. Hydrogels were further cultured in a humidified incubator with 95% air and 5% CO2 at 37 °C for subsequent experiments.

### Cell Viability Assay

Cell viability and metabolic activity were measured with alamarBlue™ HS Cell Viability Reagent (Thermo Fisher Scientific, Waltham, MA) following the manufacturer’s protocol. Briefly, cells were encapsulated into GelMA hydrogels as described above. At the time point of interest, growth medium was aspirated from the wells and new growth medium (450 µL) was added to the hydrogels, and the alamarBlue™ solution (10% final volume/50 µL) was added to each well. After 2-hour incubation on a shaker (60 rpm) in a humidified incubator with 95% air and 5% CO2 at 37 °C, the alamarBlue™ solution was measured for the fluorescence of resorufin (540 (52)-nm excitation, 580 (20-nm emission) using a F200 spectrophotometer (Tecan, Switzerland). Negative control (background) was subtracted from each measurement.

### Drug Response Assay

Drug response metrics were based on cell viability/metabolic activity assay and measured with alamarBlue™ as described above. To generate drug response curves, each of the four cell lines (8MGBA, 8MGBA-TMZres, 42MGBA, 42MGBA-TMZres) were encapsulated into GelMA hydrogels at 5 x 105 cells/mL as described above. After 24 hours of culture, initial viability (D0) was determined with alamarBlue assay as described above. After D0 measurement were collected, hydrogels were rinsed with PBS, and growth medium was supplemented with TMZ and added to the hydrogel culture as single-dose TMZ treatments ranging from 0.03 µM to 1000 µM (serial dilutions) or a DMSO vehicle control (0 μM). Without replenishing TMZ, drug response was measured at 3, 5, and 7 days after drug treatment using alamarBlue™ assay. Growth rate inhibition (GR) and related drug response metrics (GR50, GEC50, GRmax, etc.) were determined, as described in Hafner et al. (31). First, relative cell count was determined based on the metabolic activity measured with the alamarBlue™ assay. Then, GR metrics were calculated as described by Hafner et al., and the drug response metrics were calculated using the Online GR Calculator (www.grcalculator.org/grcalculator).

### Metronomic TMZ Assay

Furthermore, to evaluate the effect of metronomic (low-dose, continuous) TMZ exposure on our model of acquired TMZ resistance, cell-laden GelMA hydrogels were treated with the following sets of doses: (1) a single dose of 5 µM, or 5 repeated doses of 5 µM, or a cumulative single dose of 25 µM, and similarly, (2) a single dose of 20 µM, or 5 repeated doses of 20 µM, or a cumulative single dose of 100 µM. The repeated doses were administered in daily intervals (24 hours between each dose). Cell viability assay (as described above) was done at days 0, 3, 5, and 7 after either the single TMZ treatment for single dose exposures or after the final TMZ treatment for continual, metronomic dose exposure. Day 0 was set to be as the day of the final TMZ treatment for each experiment. Data was normalized to day 0 measurements prior to the analysis.

### Apoptosis Protein Array

GBM cells (8MGBA, 8MGBA-TMZres, 42MGBA, 42MGBA-TMZres) were cultured in GelMA hydrogels as described above for 8 days with media changes every 2-3 days. On day 1, hydrogels were treated with either (1) DMSO vehicle control or (2) TMZ at the GR50 concentration appropriate for each cell line. TMZ was not replenished afterwards. On day 8 (7 days after treatment), hydrogels were collected and washed in ice cold PBS and then frozen at -80oC. Subsequently, proteins were extracted via lysis using a lysis buffer from the Proteome Profiler™ Human Apoptosis Array Kit (ARY009, R&D Systems, Minneapolis, MN) kit. Lysate inputs were normalized by determining overall protein concentration using Pierce™ BCA Protein Assay (Thermo Fisher Scientific, Waltham, MA). Subsequently, the Proteome Profiler™ Human Apoptosis Array Kit was used following the manufacturer’s protocol. Blots were imaged using the Amersham ImageQuant 800 Fluor chemiluminescence imager (Cytiva, Marlborough, MA). Cytokine dot intensities were quantified using a MATLAB Protein Array Tool (https://www.mathworks.com/matlabcentral/fileexchange/35128-protein-array-tool). For data analysis, background (negative control/PBS) was subtracted from dot intensities, and the intensity values were further normalized to the positive control.

### Spheroid Migration Assay

Spheroids were constructed using the previously reported method (11, 23, 32). Briefly, GBM cells (8MGBA, 8MGBA-TMZres, 42MGBA, 42MGBA-TMZres) were counted and resuspended into 5000 cells/200 μL media per well and added into 96-well spheroid ultra-low attachment (ULA) microplates (Corning, Corning, NY). Plates were centrifuged at 100×g for 1 min to assist spheroid formation then placed into a humidified incubator (37oC, 5% CO2) for 24 h. Plates were then incubated for additional 24 h with gentle shaking at 60 rpm. Subsequently, formed spheroids were transferred and mixed with GelMA precursor solution and photopolymerized into hydrogels as described above. Spheroid images were acquired using a Leica DMi8 bright-field microscope (Leica, Germany) at days 0 (immediately upon seeding), 1, 2 and 3. Invasion was then quantified via ImageJ, and the invasion distance was reported as fold change of the spheroid outgrowth area compared to day 0 as described previously.

### Secretome Profiling

GBM cells (8MGBA, 8MGBA-TMZres, 42MGBA, 42MGBA-TMZres) we cultured in GelMA hydrogels as described above for 7 days. Media was changed on day 2 and 3 with partial media change on day 5. On day 7, cell culture media was collected from the hydrogels. The media (secretome) were then spun down (300×g, 10 min) to remove any debris. Subsequently, pulled secretome for each group was profiled using a Proteome Profiler™ Human Angiogenesis Array Kit (ARY007, R&D Systems, Minneapolis, MN) following the manufacturer’s protocol. Blots were imaged using the Amersham ImageQuant 800 Fluor chemiluminescence imager (Cytiva, Marlborough, MA). Cytokine dot intensities were quantified using the MATLAB Protein Array Tool (https://www.mathworks.com/matlabcentral/fileexchange/35128-protein-array-tool). For data analysis, background (negative control/PBS) was subtracted from dot intensities, and the intensity values were further normalized to positive control.

### Statistical Analysis

Statistical analysis was performed in RStudio (RStudio Inc., Boston, MA) using the programming language R (R Core Team, Vienna, Austria). Normality of the residuals for all samples was verified using the Shapiro-Wilk test, and homogeneity of variances was verified using the Levene’s test. Statistical analysis of data satisfying normality and homogeneity of variances was performed using the Student’s t-test (2 groups) or one-way analysis of variance (ANOVA) followed by Tukey honest significant difference (HSD) post hoc test (for 3+ groups). Statistical analysis of data not satisfying normality was performed using Kruskal-Wallis H test followed by Dunn’s multiple comparison post hoc test. Statistical analysis of data not satisfying homogeneity of variance was performed using Welch’s t-test/ANOVA followed by Games-Howell post hoc test. Significance for all statistical analyses was set at p < 0.05. A minimum of n = 3 samples was used for all analyses and specified in each result section. Error bars are plotted as the standard deviation (SD).

## Acknowledgments

The content herein is solely the responsibility of the authors and does not necessarily represent the official views of the National Institutes of Health. The authors also gratefully acknowledge additional funding provided by the Department of Chemical & Biomolecular Engineering and Cancer center at Illinois at the University of Illinois at Urbana-Champaign. The authors would like to thank Sydney Anne McKee (UIUC) for technical guidance that aided in experimental design and execution.

## Author contributions

We describe contributions to the manuscript using the Contributor Roles Taxonomy (CRediT) (76, 77):

Conceptualization: VK, BACH, RBR

Methodology: VK

Investigation: VK, EE

Visualization: VK

Formal Analysis: VK

Data Curation: VK

Project Administration: BACH

Supervision: BACH, RBR

Writing—original draft: VK

Writing—review & editing: VK, BACH, RBR

Resources: BACH

Funding Acquisition: BACH

## Data and materials availability

All data needed to evaluate the conclusions in the paper are present in the paper and the Supplementary Materials.

