## Supplemental Materials for "Acquired temozolomide resistance instructs patterns of glioblastoma behavior in gelatin hydrogels"

Victoria Kriuchkovskaia *et al.*

*Brendan A.C. Harley.

**This PDF file includes:**

Figs. S1 to S8
Tables S1 to S4

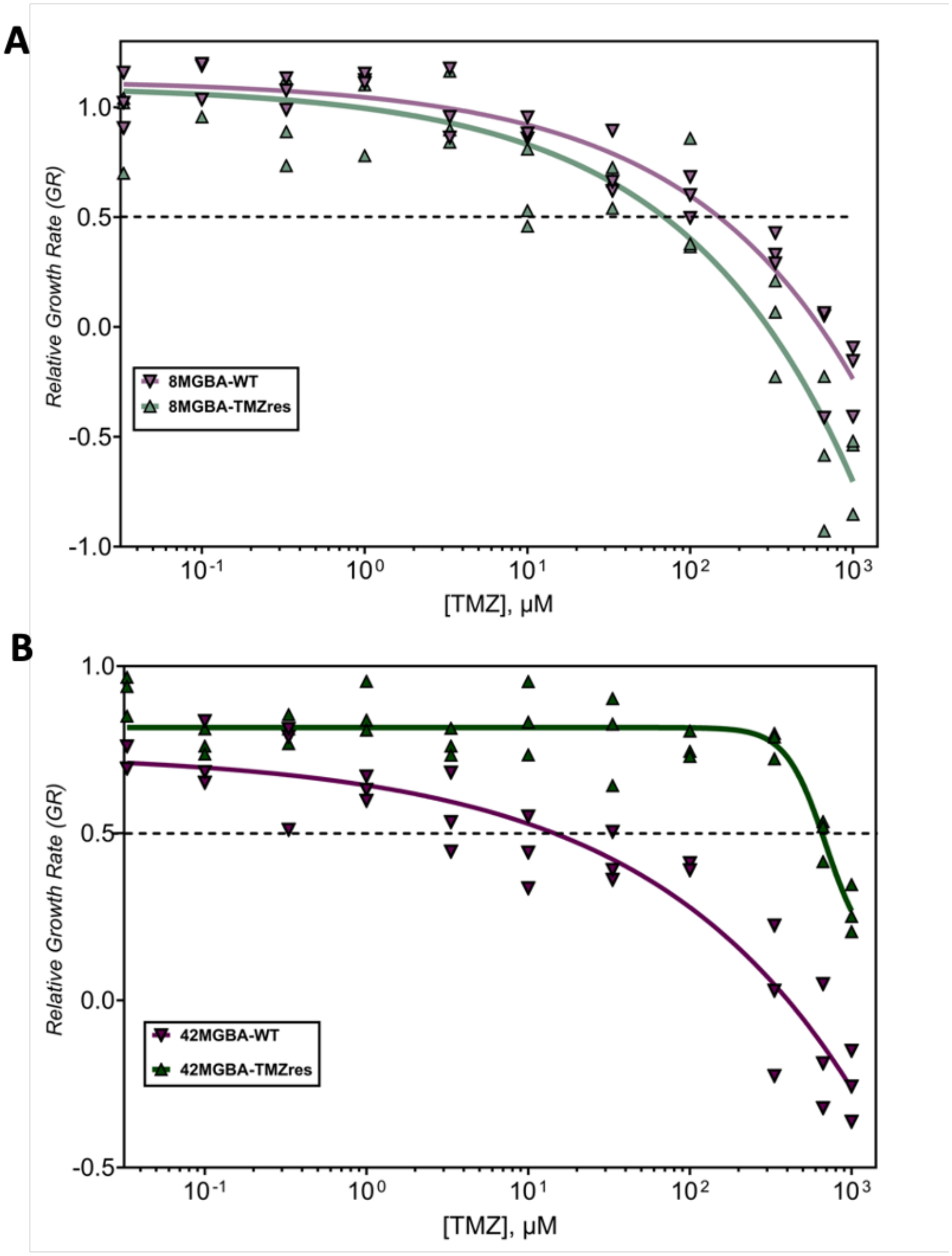


**Fig. S1.**

**Dose-response curves of TMZ-sensitive vs. TMZ-resistant glioblastoma cell lines cultured in GelMA hydrogels 3 days after TMZ treatment. (A)** Dose response, growth-rate inhibition curves of the 8MGBA vs. 8MGBA-TMZres cell lines 3 days after TMZ treatment. **(B)** Dose response, growth-rate inhibition curves of the 42MGBA vs. 42MGBA-TMZres cell lines 3 days after TMZ treatment. Each individual data point along with fitted GR curves are shown. N = 3 hydrogels per condition.

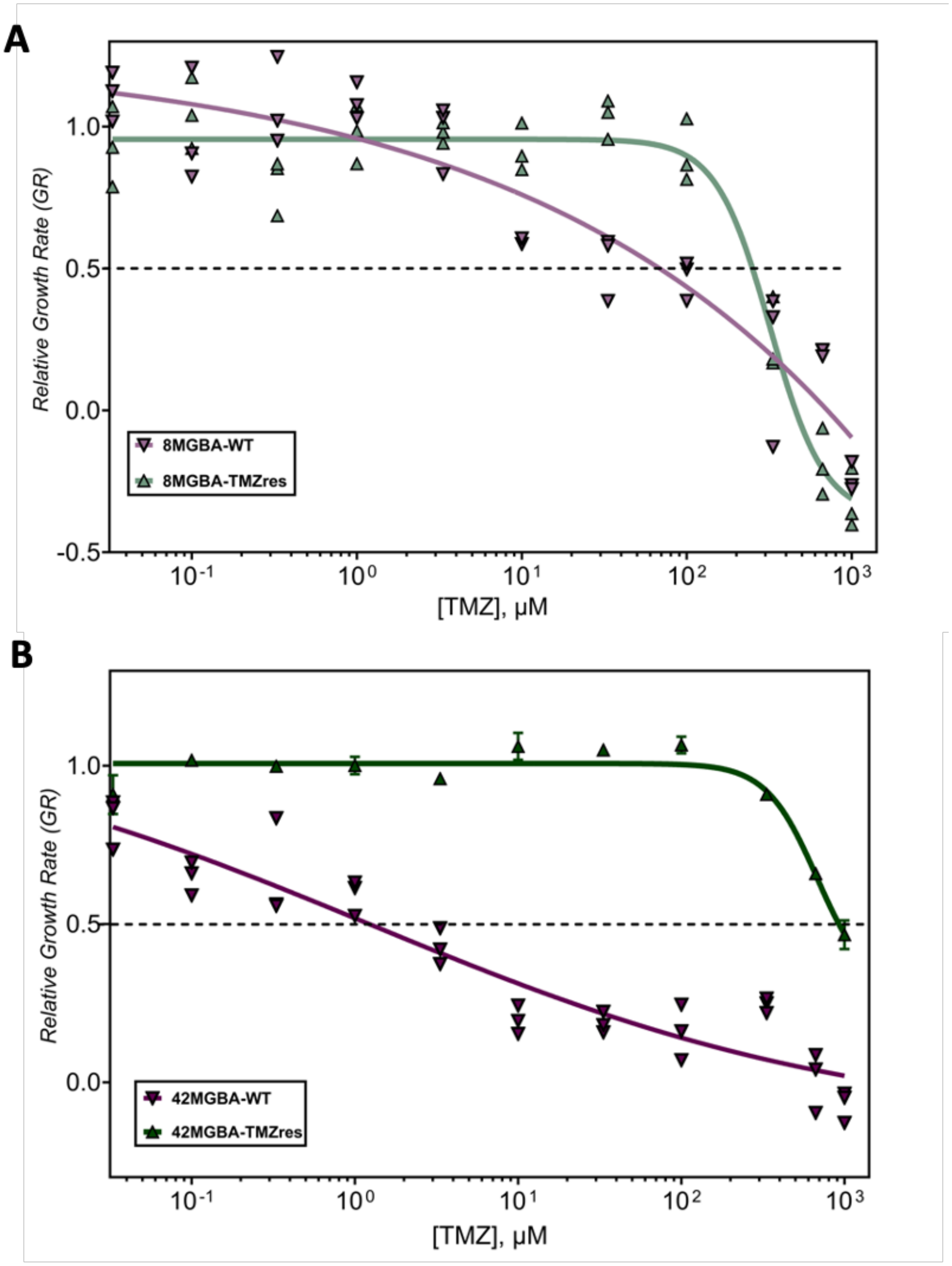


**Fig. S2.**

**Dose-response curves of TMZ-sensitive vs. TMZ-resistant glioblastoma cell lines cultured in GelMA hydrogels 5 days after TMZ treatment. (A)** Dose response, growth-rate inhibition curves of the 8MGBA vs. 8MGBA-TMZres cell lines 5 days after TMZ treatment. **(B)** Dose response, growth-rate inhibition curves of the 42MGBA vs. 42MGBA-TMZres cell lines 5 days after TMZ treatment. Each individual data point along with fitted GR curves are shown. N = 3 hydrogels per condition.

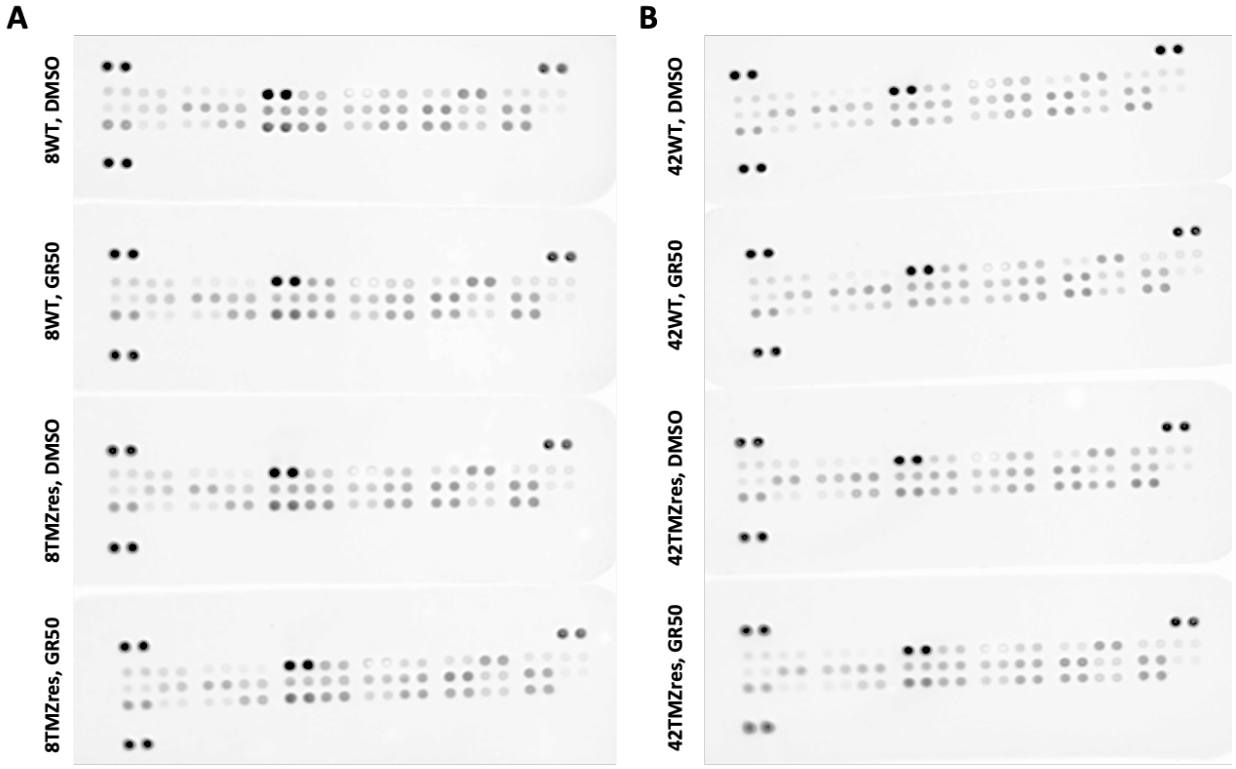


**Fig. S3.**

**Scanned images of the Proteome Profiler™ Human Apoptosis Array Kit blots. (A)** Blots of 8MGBA-WT/TMZres samples treated either with DMSO vehicle control or TMZ at GR50 doses. **(B)** Blots of 42MGBA-WT/TMZres samples treated either with DMSO vehicle control or TMZ at GR50 doses.

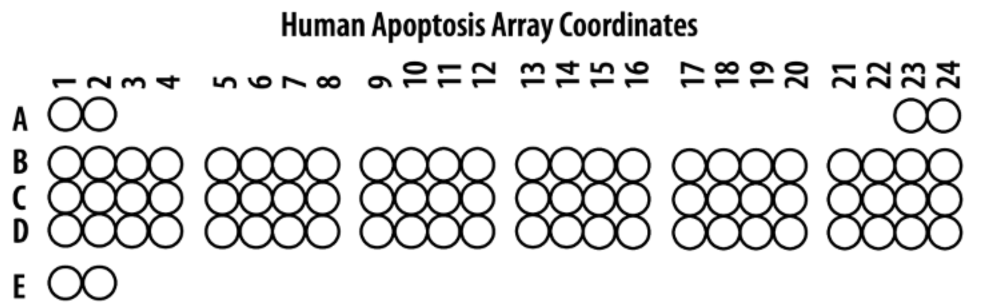


| Coordinate | Target/Control | Coodinate | Target/Control |
|---|---|---|---|
| *A1, A2* | Positive Control | *C13, C14* | HO-2/HMOX2 |
| *A23, A24* | Positive Control | *C15, C16* | HSP27 |
| *B1, B2* | BAD | *C17, C18* | HSP60 |
| *B3, B4* | BAX | *C19, C20* | HSP70 |
| *B5, B6* | BCL2 | *C21, C22* | HTRA2/Omi |
| *B7, B8* | BCLX | *C23, C24* | Livin |
| *B9, B10* | Pro-Caspase-3 | *D1, D2* | PON2 |
| *B11, B12* | Cleaved Caspase-3 | *D3, D4* | p21/CIP1/CDKN1A |
| *B13, B14* | Catalase/CAT | *D5, D6* | p27/Kip1 |
| *B15, B16* | cIAP-1/BIRC2 | *D7, D8* | Phospho-p53 (S15) |
| *B17, B18* | cIAP-2/BIRC3 | *D9, D10* | Phospho-p53 (S46) |
| *B19, B20* | Claspin/CLSPN | *D11, D12* | Phospho-p53 (S392) |
| *B21, B22* | Clusterin/CLU | *D13, D14* | Phospho-Rad17 (S635) |
| *B23, B24* | Cytochrome C/CYCS | *D15,D16* | SMAC/Diablo |
| *C1, C2* | TRAIL R1/DR4 | *D17,D18* | Survivin/BIRC5 |
| *C3, C4* | TRAIL R2/DRS | *D19, D20* | TNF RI/TNFRSF1A |
| *C5, C6* | FADD | *D21, D22* | XIAP |
| *C7, C8* | Fas/TNFRSF6/CD95 | *D23, D24* | PBS (Negative Control) |
| *C9, C10* | HIF-1a | *E1, E2* | Postivie Control |
| *C11, C12* | HO-1/HMOX1/HSP32 | | |

**Fig. S4.**

**Coordinates the Proteome Profiler™ Human Apoptosis Array Kit blots used to analyze data.**

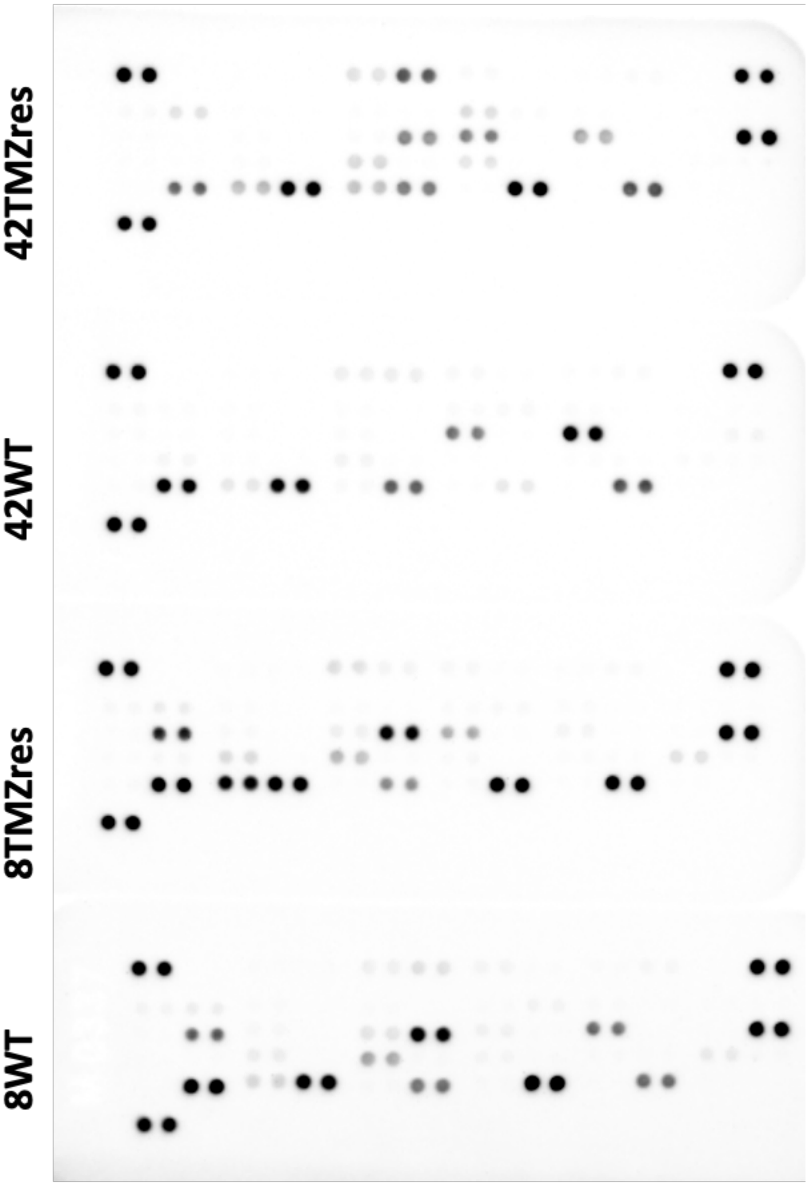


**Fig. S5.**
**Scanned images of the Proteome Profiler™ Human Angiogenesis Array Kit blots.**

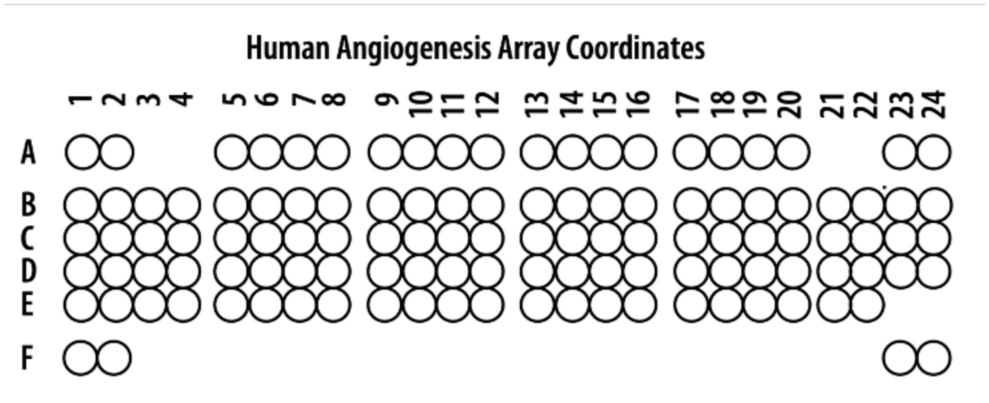


| Coordinate | Target/Control | Coodinate | Target/Control |
|---|---|---|---|
| *A1, A2* | Reference Spots | *C17, C18* | IL-8 |
| *A5, A6* | Activin A | *C19, C20* | LAP (TGF-B1) |
| *A7, A8* | ADAMTS-1 | *C21, C22* | Leptin |
| *A9, A10* | Angiogenin | *C23, C24* | MCP-1 |
| *A11, A12* | Angiopoietin-1 | *D1, D2* | MIP-1a |
| *A13, A14* | Angiopoietin-2 | *D3, D4* | MMP-8 |
| *A15, A16* | Angiostatin/Plasminogen | *D5, D6* | MMP-9 |
| *A17, A18* | Amphiregulin | *D7, D8* | NRG1-B1 |
| *A19, A20* | Artemin | *D9, D10* | Pentraxin 3 (PTX3) |
| *A23, A24* | Reference Spots | *D11, D12* | PD-ECGF |
| *B1, B2* | Coagulation Factor III | *D13, D14* | PDGF-AA |
| *B3, B4* | CXCL16 | *D15, D16* | PDGF-AB/PDGF-BB |
| *B5, B6* | DPPIV | *D17, D18* | Persephin |
| *B7, B8* | EGF | *D19, D20* | Platelet Factor 4 (PF4) |
| *B9, B10* | EG-VEGF | *D21, D22* | PIGF |
| *B11, B12* | Endoglin | *D23, D24* | Prolactin |
| *B13, B14* | Endostatin/Collagen XVIII | *E1, E2* | Serpin B5 |
| *B15, B16* | Endothelin-1 | *E3, E4* | Serpin E1 |
| *B17, B18* | FGF acidic | *E5, E6* | Serpin F1 |
| *B19, B20* | FGF basic | *E7, E8* | TIMP-1 |
| B21, B22 | FGF-4 | *E9, E10* | TIMP-4 |
| B23, B24 | FGF-7 | *E11, E12* | Thrombospondin-1 |
| C1, C2 | GDNF | *E13, E14* | Thrombospondin-2 |
| C3, C4 | GM-CSF | *E15, E16* | uPA |
| C5, C6 | HB-EGF | *E17, E18* | Vasohibin |
| C7, C8 | HGF | *E19, E20* | VEGF |
| C9, C10 | IGFBP-1 | *E21, E22* | VEGF-C |
| C11, C12 | IGFBP-2 | *F1, F2* | Reference Spots |
| C13, C14 | IGFBP-3 | *F23, F24* | Negative Control |
| C15, C16 | IL-1B | | |

**Fig. S6.**

**Coordinates the Proteome Profiler™ Human Angiogenesis Array Kit blots used to analyze data.**

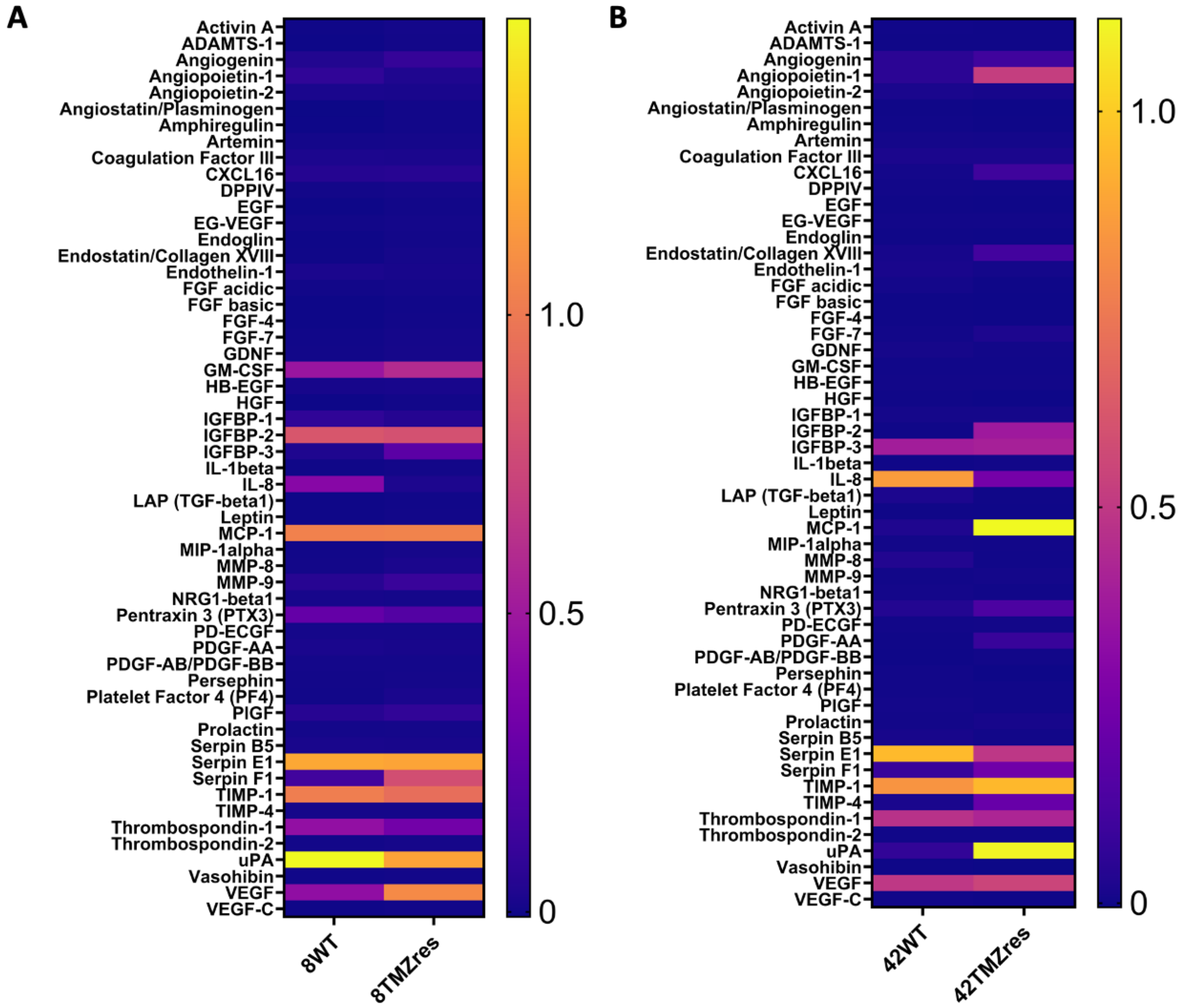


**Fig. S7.**

**Expression of matrix remodeling and angiogenesis-related cytokines. (A)** Heat map showing the normalized relative expression of all profiled matrix remodeling and angiogenesis-related cytokines in the secretome collected from 8MGBA-WT vs. TMZ-res cells after 7 days of culture in GelMA hydrogels. **(B)** Heat map showing the normalized relative expression of all profiled matrix remodeling and angiogenesis-related cytokines in the secretome collected from 42MGBA-WT vs. TMZ-res cells after 7 days of culture in GelMA hydrogels

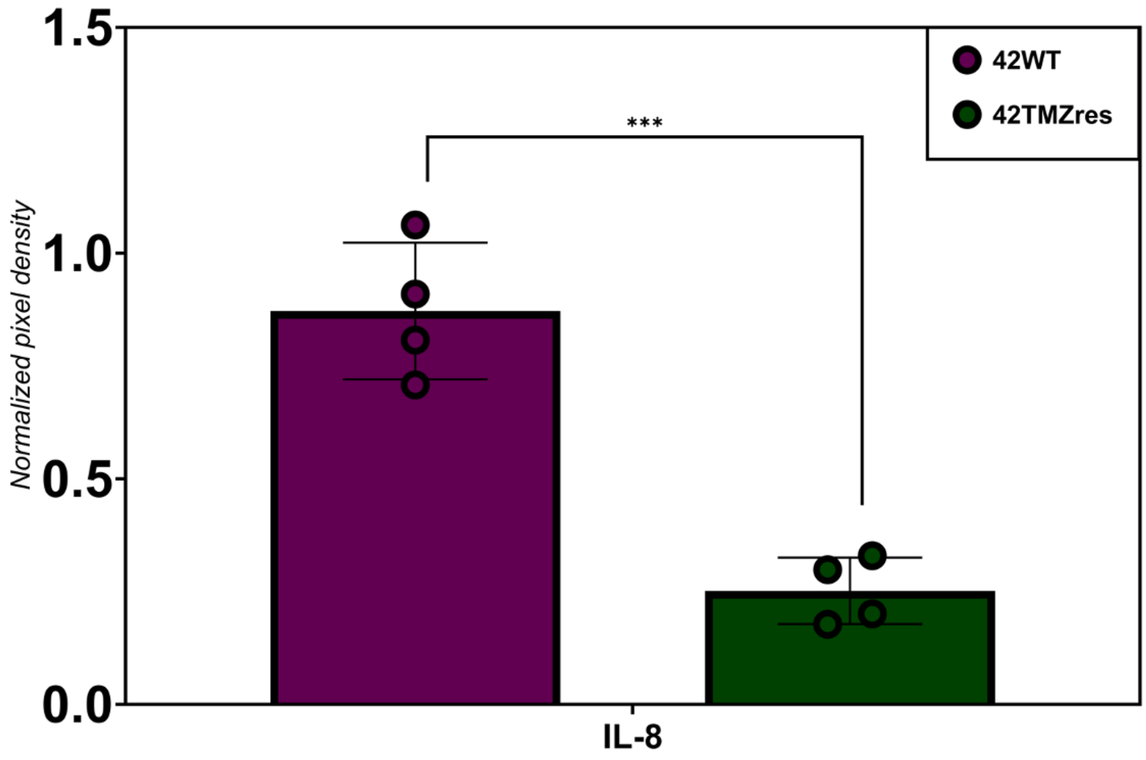


**Fig. S8.**

Relative expression levels of IL-8 in the 42MGBA pair.

**Table S1.**
**Results of statistical analysis of expression data of apoptosis-related proteins in the 8MGBA pair.**

| *Groups: A-8WT DMSO, B-8WT TMZ, C-8TMZres DMSO, D-8TMZres TMZ* | | | |
|---|---|---|---|
| *8MGBA* | *Significance* | *p-value* | *post hoc* |
| **BAD** | | 3.20E-04 | A-C, B-C, C-D |
| **BAX** | | 6.94E-02 | |
| **BCL2** | | 6.53E-03 | A-D |
| **BCLX** | | 5.32E-03 | A-C, A-D |
| **Pro-CASP3** | | 1.88E-05 | A-B, A-C, A-D, B-C, C-D |
| **Cleaved CASP3** | | 6.20E-03 | A-B, A-D, B-C, C-D |
| **CAT** | | 6.41E-01 | |
| **BIRC2** | | 9.73E-03 | A-B, A-D, B-C, C-D |
| **BIRC3** | | 1.08E-01 | |
| **CLSPN** | | 4.49E-07 | A-D, B-C, B-D, C-D |
| **CLU** | | 5.76E-02 | |
| **CYCS** | | 3.12E-06 | A-D, B-D, C-D |
| **TRAIL-R1** | | 3.27E-03 | A-D, B-C, B-D |
| **TRAIL-R2** | | 5.23E-06 | A-B, A-C, A-D |
| **FADD** | | 4.50E-02 | A-D |
| **FAS** | | 1.21E-07 | A-C, A-D, B-C, B-D, C-D |
| **HIF1A** | | 3.88E-06 | A-B, A-C, A-D, B-C, C-D |
| **HMOX1** | | 4.77E-04 | A-B, A-D, C-D |
| **HMOX2** | | 7.36E-02 | |
| **HSP27** | | 7.36E-01 | |
| **HSP60** | | 1.82E-07 | |
| **HSP70** | | 3.62E-02 | A-C |
| **HTRA2** | | 3.21E-05 | A-C, B-C, B-D, C-D |
| **Livin** | | 1.42E-04 | A-D, B-D, C-D |
| **PON2** | | 6.95E-02 | |
| **CDNK1A** | | 7.85E-05 | A-B, A-C, A-D |
| **Kip1** | | 3.45E-03 | A-B, A-C, A-D |
| **Phospho-p53 (S15)** | | 5.48E-08 | A-B, A-C, A-D, B-D, C-D |
| **Phospho-p53 (S46)** | | 1.98E-03 | A-B, A-D, B-C |
| **Phospho-p53 (S392)** | | 1.69E-05 | A-B, A-D, B-C, B-D, C-D |
| **Phospho-Rad17 (S635)** | | 9.57E-05 | A-D, B-C, B-D, C-D |
| **DIABLO** | | 6.40E-03 | A-D, B-C, B-D |
| **BIRC5** | | 1.13E-11 | A-B, A-D, B-C, C-D |
| **TNFRSF1A** | | 5.71E-08 | A-B, A-C, A-D, B-C, B-D, C-D |
| **XIAP** | | 1.21E-08 | A-C, A-D, B-C, B-D, C-D |

**Table S2.**

**Results of statistical analysis of expression data of apoptosis-related proteins in the 42MGBA pair.**

| *Groups: A-42WT DMSO, B-42WT TMZ, C-42TMZres DMSO, D-42TMZres TMZ* | | | |
|---|---|---|---|
| ***42MGBA*** | ***Significance*** | ***p-value*** | ***post hoc*** |
| **BAD** | | 3.23E-04 | A-B, A-C, A-D |
| **BAX** | | 1.86E-01 | |
| **BCL2** | | 4.18E-01 | |
| **BCLX** | | 1.67E-02 | A-C |
| **Pro-CASP3** | | 4.55E-07 | A-B, A-C, A-D, B-D, C-D |
| **Cleaved CASP3** | | 3.95E-07 | A-B, A-D, B-C, C-D |
| **CAT** | | 9.47E-03 | A-C, C-D |
| **BIRC2** | | 2.40E-03 | A-C, C-D |
| **BIRC3** | | 5.22E-08 | A-C, A-D, B-C, B-D, C-D |
| **CLSPN** | | 2.17E-08 | A-B, A-C, A-D, B-D, C-D |
| **CLU** | | 1.89E-15 | A-C, A-D, B-C, B-D, C-D |
| **CYCS** | | 5.37E-02 | |
| **TRAIL-R1** | | 5.22E-04 | A-C, B-C, C-D |
| **TRAIL-R2** | | 6.27E-07 | A-C, A-D, B-C, B-D, C-D |
| **FADD** | | 1.08E-04 | A-D, B-C, B-D |
| **FAS** | | 3.60E-11 | A-B, A-C, A-D, B-D, C-D |
| **HIF1A** | | 5.23E-08 | A-B, A-C, A-D, B-D, C-D |
| **HMOX1** | | 2.35E-09 | A-C, A-D, B-C, B-D |
| **HMOX2** | | 8.81E-10 | A-B, A-C, A-D, B-D, C-D |
| **HSP27** | | 2.57E-05 | A-C, A-D, B-C, B-D |
| **HSP60** | | 3.81E-08 | A-B, A-D, B-C, B-D, C-D |
| **HSP70** | | 5.48E-03 | B-C |
| **HTRA2** | | 3.31E-10 | A-D, B-D, C-D |
| **Livin** | | 3.21E-03 | A-C, A-D |
| **PON2** | | 3.33E-05 | A-B, A-C, B-D, C-D |
| **CDNK1A** | | 3.52E-01 | |
| **Kip1** | | 1.30E-01 | |
| **Phospho-p53 (S15)** | | 1.51E-07 | A-C, A-D, B-C, B-D, C-D |
| **Phospho-p53 (S46)** | | 2.16E-14 | A-C, A-D, B-C, B-D, C-D |
| **Phospho-p53 (S392)** | | 1.01E-13 | A-C, A-D, B-C, B-D, C-D |
| **Phospho-Rad17 (S635)** | | 6.66E-15 | A-B, A-C, A-D, B-C, B-D, C-D |
| **DIABLO** | | 1.92E-09 | A-B, A-C, A-D, B-C, B-D, C-D |
| **BIRC5** | | 2.30E-06 | A-B, A-C, B-C, B-D, C-D |
| **TNFRSF1A** | | 2.10E-13 | A-B, A-C, A-D, B-C, B-D, C-D |
| **XIAP** | | 6.29E-10 | A-C, A-D, B-C, B-D, C-D |

**Table S3.**

**Results of statistical analysis of expression data of angiogenesis-related cytokines in the 8MGBA and 42MGBA pairs.**

| *Groups: 8WT vs 8TMZres* | *Significance* | *p-value* | *Test* |
|---|---|---|---|
| **ANG** | | 8.65E-02 | *t-test* |
| **GM-CSF** | | 3.35E-01 | *Welch* |
| **IGFBP-2** | | 7.40E-01 | *t-test* |
| **IGFBP-3** | | 1.06E-02 | *Welch* |
| **IL-8** | | 6.75E-03 | *Welch* |
| **MCP-1** | | 9.65E-01 | *t-test* |
| **MMP-9** | | 1.13E-01 | *Welch* |
| **PTX3** | | 3.61E-01 | *t-test* |
| **PlGF** | | 2.16E-01 | *t-test* |
| **Serpin E1** | | 9.23E-01 | *Welch* |
| **Serpin F1** | | 2.20E-03 | *Welch* |
| **TIMP-1** | | 6.43E-01 | *Welch* |
| **THBS1** | | 2.23E-01 | *Welch* |
| **uPA/PLAU** | | 1.52E-01 | *Welch* |
| **VEGF** | | 2.26E-03 | *Welch* |
| *Groups: 42WT vs 42TMZres* | *Significance* | *p-value* | *Test* |
| **ANG** | | 1.31E-01 | *Welch* |
| **ANGPT1** | | 2.62E-03 | *Welch* |
| **CXCL16** | | 3.09E-02 | *Welch* |
| **Endostatin** | | 2.54E-02 | *Welch* |
| **IGFBP-2** | | 6.34E-03 | *Welch* |
| **IGFBP-3** | | 9.48E-01 | *t-test* |
| **IL-8** | | 3.21E-04 | *t-test* |
| **MCP-1** | | 1.43E-03 | *Welch* |
| **PTX3** | | 3.49E-02 | *Welch* |
| **PDGFA** | | 2.28E-02 | *Welch* |
| **Serpin E1** | | 1.48E-02 | *Welch* |
| **Serpin F1** | | 3.54E-02 | *Welch* |
| **TIMP-1** | | 4.09E-01 | *t-test* |
| **TIMP-4** | | 1.71E-02 | *Welch* |
| **THBS1** | | 5.37E-01 | *t-test* |
| **uPA/PLAU** | | 1.62E-03 | *Welch* |
| **VEGF** | | 4.75E-01 | *t-test* |
